# Quantitative multi-scale morphodynamic analysis reveals ratchet-like collective DVE migration and epiblast retrograde cell flow during anterior patterning in the mouse embryo

**DOI:** 10.64898/2026.04.24.720339

**Authors:** Matthew J. Stower, Felix Y. Zhou, Rahil Valani, Jan Rozman, Holly Hathrell, Jonathan Godwin, Xin Lu, Jens Rittscher, Julia M. Yeomans, Shankar Srinivas

## Abstract

The collective unidirectional migration of distal visceral endoderm (DVE) cells in the early mouse embryo is required to pattern the anterior-posterior (A-P) axis in the epiblast. It is unknown to what extent A-P axial asymmetries exist prior to DVE migration, how migration becomes channeled towards one side of the embryo, and whether the epiblast cells they migrate over have coordinated movements. We developed a quantitative embryo-wide, tissue-tracking approach to analyse visceral endoderm and epiblast tissue morphodynamics in a longitudinal light-sheet imaged, multi-embryo data-set. Here we show that asymmetric morphology of the ectoplacental cone already present prior to DVE migration correlates with the alignment of the A-P axis but not its polarity. DVE cell movements are initiated with a relatively low cell coordination and small net migration, then get channelled in an abrupt transition to a highly coordinated, uni-directional anterior motion. This anteriorwards migration is characterised by a ratchet-like, intermittent motion. Vertex modelling demonstrates that tissue rheology can account for DVE start–stop motion, and suggests that T1-mediated stress relaxation in the surrounding tissue can facilitate intermittent DVE motion without requiring intrinsic fluctuations in DVE velocity. Finally, comparing cell movement of the DVE with the underlying epiblast reveals a previously unknown coordinated motion in the anterior epiblast, opposite to the direction of DVE migration. Together these data provide insights into the origin of embryonic axial asymmetry and a previously unappreciated coordination between VE and epiblast tissue-motion during anterior-posterior patterning.

**GRAPHICAL ABSTRACT:** 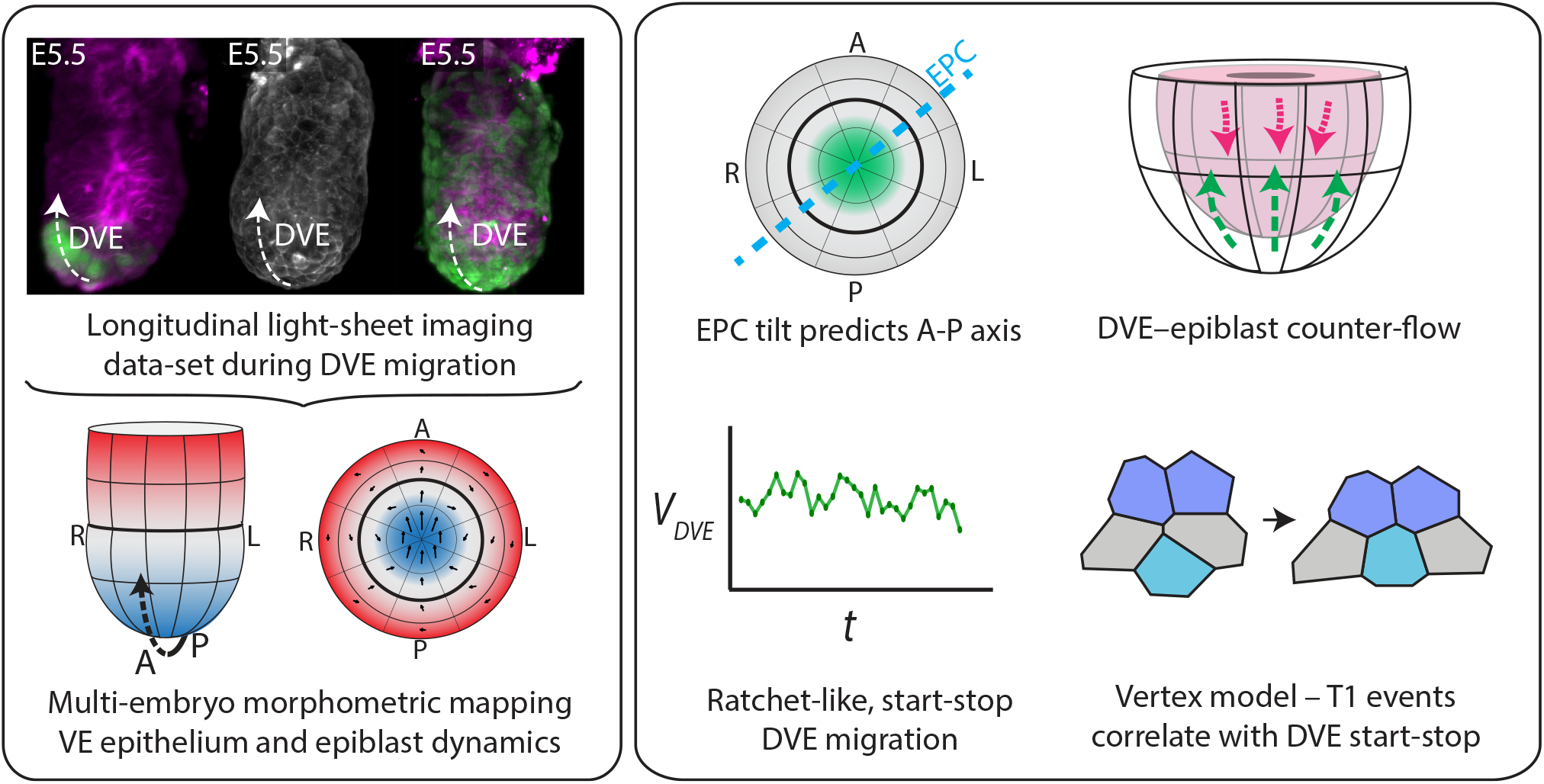

See supplemental movie: animated abstract

## INTRODUCTION

In embryonic day 5.5 (E5.5) mouse embryos, the epiblast that will form the foetus is arranged as a pseudostratified epithelium surrounded by a monolayer epithelium, the visceral endoderm (VE) (Figure 1A). At E5.5 a sub-population of VE cells located in the distal-most region of the tissue (distal visceral endoderm – DVE) are induced and undergo a collective migration towards one side of the embryo (Srinivas et al. 2004; P. Q. Thomas et al. 1998). Secretion of inhibitors of the TGF-β/NODAL (Yamamoto et al. 2004) and WNT (Belo et al. 1997) pathways by these cells (at the future anterior) results in the restriction of the primitive streak, the site of gastrulation, to the opposite (posterior) side of the epiblast by E6.25. Interventions affecting the induction or migration of DVE cells leads to an incorrectly patterned, non-viable embryo (P. Thomas and Beddington 1996), demonstrating the critical role of this tissue in embryonic development.

**Figure 1.**
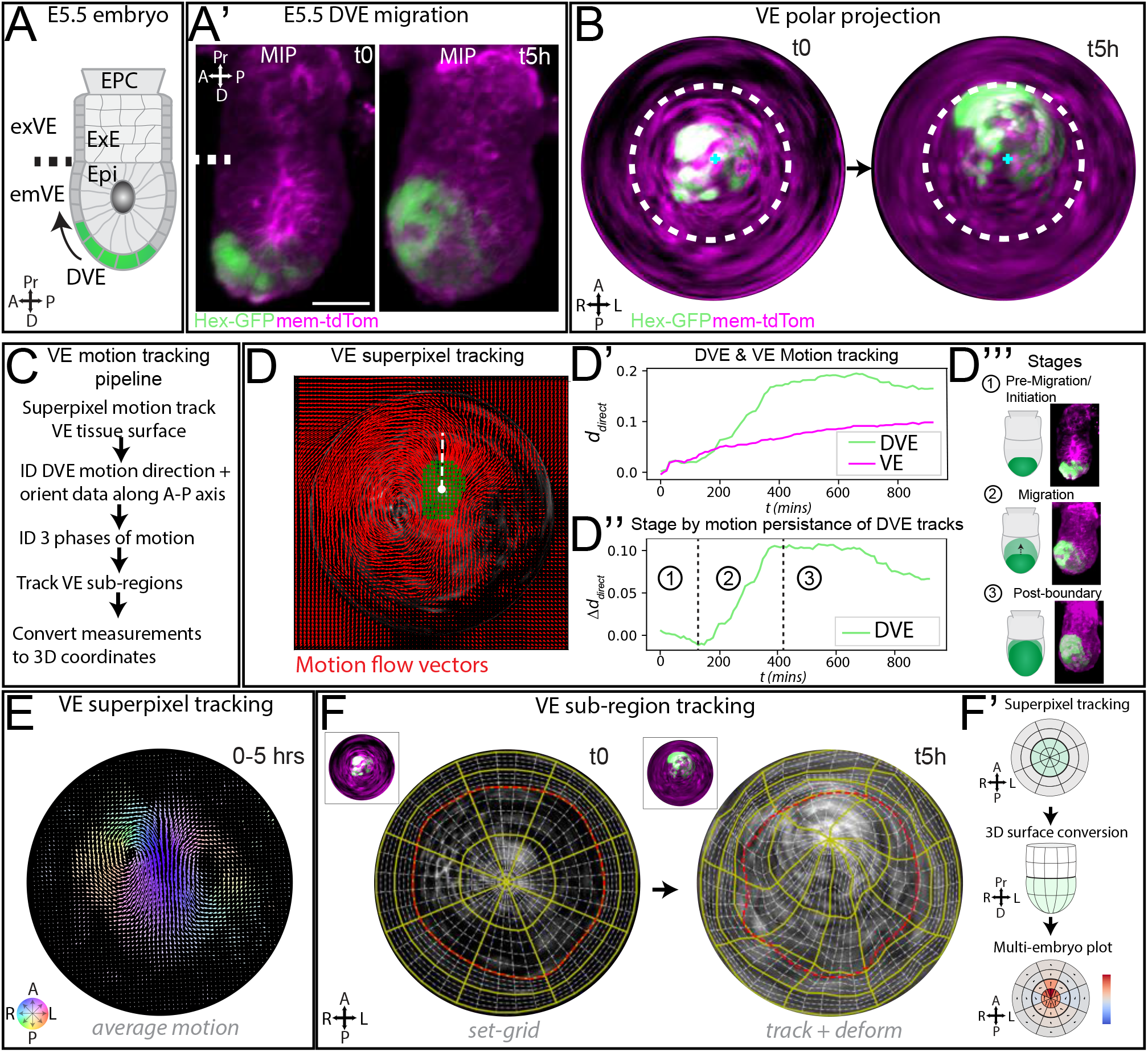
Data processing pipeline for superpixel motion-based analysis of VE tissue morphodynamics. **(A)** Diagram of E5.5 mouse embryo; Epi = epiblast, ExE = extra-embryonic ectoderm, EPC = ectoplacental cone, emVE = embryonic visceral endoderm, exVE = extra-embryonic visceral endoderm, DVE = distal visceral endoderm. **(A’)** Lateral view of example E5.5 Hex-GFP:mem-tdTomato embryo at two time-points, imaged using a ZEISS Z.1 light-sheet at a 10-minute interval. MIP = max intensity projection. Scale bar = 50 μm. **(B)** Polar geodesic projection of the apical VE surface of embryo in A’ . Re-projections enable visualisation of entire VE and the application of sophisticated motion analysis tools to analyse time-lapse data. Dotted line shows position of emVE-exVE boundary – the limit of DVE cell migration. Blue cross = distal tip. **(C)** Overview of VE motion-tracking pipeline for quantitative analysis of time-lapse data-set. **(D’-D’’)** Automatic staging of DVE migration from superpixel-based motion-tracking. **(D)** VE superpixel motion tracking on VE polar projection; red arrows show motion-flow vectors. DVE-core motion highlighted in green. Dotted line shows mean consensus angle vector of DVE which is used to align embryos along their A-P axis. The persistence of motion direction *d*_*direct*_ of DVE was separated from the surrounding VE. **(D’’)** The motion persistence of DVE showed three phases; 1) pre-migration/initiation, 2) migration to boundary, 3) post-boundary, as shown in the max intensity projections. **(D’’’)** Example embryo showing last timepoint of each phase. **(E)** A flow-like pattern of motion across the VE was observed by superpixel motion tracking. Colours denote mean vector angle. **(F)** To track sub-regions of the VE, a deformable grid was seeded at the start of each phase. By tracking the motion within each sub-region the grid becomes deformed and multiple parameters are analysed. **(F’)** To quantify parameters and to combine data from across embryos, 2D tracking data is converted back to 3D coordinate-space for analysis which can be summarised in a multi-embryo polar plot. Note re-projections have no scale bar as they are non-linear.

DVE cell migration is an active process, as first shown by live imaging of the Hex-GFP mouse line in which DVE cells are labelled with GFP (Srinivas et al. 2004). This revealed that the DVE has basally-localised cell projections polarised in the direction of migration (Migeotte et al. 2010; Srinivas et al. 2004). In the absence of proteins such as RAC1, or components of the WAVE complex that regulate the cytoskeleton, migration is aberrant or inhibited (Bloomekatz et al. 2012; Migeotte et al. 2010; Omelchenko et al. 2014; Rakeman and Anderson 2006). Throughout DVE migration the VE remains as an epithelial monolayer with intact tight-junctions and adherens-junctions (Trichas et al. 2011), meaning DVE cells negotiate their way through surrounding VE cells that must accommodate their movement. We have recently quantitatively assessed, at single-cell resolution, the component cellular events during DVE migration (cell movements, shape changes, rearrangements, divisions, etc.). This has revealed that spatial and temporal heterogeneity in cell behaviour, morphology and mechanical tension facilitate, as well as set the bounds to, DVE migration (Stower et al. 2023).

However, what determines the *direction* of DVE migration remains unclear. Several asymmetries have been described that could influence migration direction. Molecular asymmetries include the expression pattern of *Lefty1* and *Cer1* in the forming primitive endoderm of the pre-implantation blastocyst that will give rise to the DVE (Takaoka et al. 2006; Takaoka et al. 2011; Torres-Padilla et al. 2007), where they remain asymmetrically expressed prior to DVE migration (Takaoka et al. 2011; Thowfeequ et al. 2024; Yamamoto et al. 2004), as well as the reported dynamic expression of OTX-2 in the emVE (Hoshino et al. 2015). We have shown that the emVE anterior-proximal to the DVE undergoes cell shape changes and cell-cell intercalation events after the onset of DVE migration (Stower et al. 2023), but it remains unclear whether there are pre-patterned *behavioural* asymmetries in the emVE.

Morphological asymmetries that could bias the direction of migration include the ‘tilt’ of the Inner Cell Mass with respect to the proximal-distal axis of the blastocyst, which renders the pre-implantation blastocyst bilaterally (rather than radially) symmetrical (Beddington and Robertson 1999). This results in the forming primitive endoderm (from which the VE is derived) also being tilted with respect to the proximal-distal axis of the blastocyst, which could lead to an offset in the position of the forming DVE with respect the distal tip, thereby biasing its migration to one side. Due to this offset the ectoplacental cone (EPC) can also form askew. It was initially shown in fixed embryos at E6.5 that the EPC aligns with the A-P axis, even though it does not predict the *polarity* of this axis (Gardner et al. 1992). This finding has recently been supported by analysis of a stage-series of E5.5 embryos (Hadjikypri et al. 2024). However, these studies have analysed fixed embryos, during stages when the embryo undergoes substantial changes in global shape (Mesnard et al. 2004; Perea-Gomez et al. 2004) and size (Snow 1977). A longitudinal study in individually tracked embryos is required to understand whether EPC morphology correlates with the direction of DVE migration, and whether such asymmetries exist *prior* to directional DVE migration.

The DVE migrates from the distal tip of the embryo over the epiblast. In mouse embryos, pharmacological inhibition of cell proliferation in the epiblast prevents DVE migration (Stuckey et al. 2011), suggesting that the growth of this tissue influences events in the overlying VE. Nodal null embryonic stem cells have also been shown to contribute preferentially to the anterior region of the epiblast in chimaeric embryos, suggesting that epiblast cell movements may not be random (Lu and Robertson 2004). Coordinated cell movements in the epiblast have been observed prior to gastrulation in the other vertebrates (Cui et al. 2005; Graper 1929; Halacheva et al. 2011). Recently, time-lapse studies have shown that in quail embryos, there is coordinated movement between the hypoblast and epiblast, prior to gastrulation (Villedieu et al. 2025). While live imaging of the E5.5 mouse epiblast has been performed (Ichikawa et al. 2013), it is unclear how coordinated any movements are with respect to the onset, or direction, of DVE migration.

To understand how multiple tissue inputs control the faithful formation of the A-P axis in the E5.5 mouse embryo, we previously generated and analysed a multi-embryo, longitudinal dataset of fluorescent reporter embryos that captures events prior to and through DVE migration at high temporal and spatial resolution (Stower et al. 2023). We also developed a computational analysis pipeline to enable the digital extraction and re-projection of both the apical surface of the VE and the basal surface of the epiblast. Here we expanded our dataset by time-lapse live imaging of dual-colour embryos expressing eGFP in the VE and membrane-targeted tdTomato in the epiblast. We then further developed a suite of tools using the MOtion SEnsing Superpixels (MOSES) (Zhou et al. 2019) framework that enabled detailed spatiotemporal analysis of VE cell motion to be integrated with multi-scale analysis of the morphology of the whole embryo. We find that EPC tilt aligns with the anterior-posterior axis at E5.5, prior to DVE migration. Initiation of DVE cell migration is characterised by a low level of coordination amongst component cells with low net migration, followed by a transition to increased directional coordination and high net migration. As cells proceed anteriorly they do so with a start-stop ratchet-like motion. An *in silico* vertex model of the VE shows that DVE start–stop motion coincides with T1 transitions in surrounding cells, a correlation also observed *in vivo*. Together, these results suggest that ratchet-like DVE motion arises from a balance between active DVE migration and mechanical resistance from the emVE, with stress relieved through T1-mediated cell rearrangements. Significantly, we also show a previously unknown coordinated motion in the anterior epiblast, opposite to the direction of DVE migration, indicating that the role of the DVE in anterior patterning is more complex than currently appreciated.

## RESULTS

### Analysis pipeline for multi-tissue analysis across scales during DVE migration

To understand if there are morphological or behavioural asymmetries in the E5.5 mouse embryo (Figure 1A) that underlie A-P axis formation requires an *in toto* live data-set to capture the 3D temporal dynamics of embryonic (epiblast) and extra embryonic (VE and EPC) tissues, as well as a quantitative analysis framework to compare different tissues within embryos and across samples. We have previously generated a multi-embryo dataset using a ZEISS Z.1 light-sheet microscope to capture 3D volumes of fluorescent reporter mouse embryos (Stower et al. 2023). This included Hex-GFP:membrane-tdTomato embryos where DVE cells are labelled by cytoplasmic eGFP in a background where all cells are labelled by membrane-targeted tdTomato (n = 9) (Figure 1A’, Movie S1) and Lifeact-GFP embryos where F-actin is ubiquitously labelled (n = 9) (Stower et al. 2023) (Figure S1B, Movie S2). We also developed a computational analysis pipeline ‘STrEAMs’ (Stower et al. 2023) to spatiotemporally register each time-lapse volume dataset and reproject VE surface coordinates as a series of 2D geodesic cartographic projections (Figure 1B, Figure S1A-B). The pipeline objectively aligns embryos along their anterior-posterior (A-P) axis using the persistent direction of DVE migration, enabling any angle measurements (e.g., the morphological tilt of the EPC) to be made with a consistent reference. It also provides a spatiotemporally consistent 3D-to-2D coordinate framework for each embryo, so that any features observed in the 2D representations can be converted back to 3D coordinates to obtain accurate measurements (Figure 1C).

To quantitatively assess local and global tissue-level behaviours, we used MOSES (Zhou et al. 2019) (Figure 1D, Figure S2). We applied our previously developed motion classifier to all time-lapse data to stage each embryo into three phases: Phase I: ‘pre-migration/initiation (where there is no persistent directional movement, i.e. no cells show unidirectional movement extending beyond 3 frames (=15 minutes)), Phase II: ‘migration’ (directional movement from distal-tip to emVE-exVE boundary) and Phase III:’post-boundary migration’ (phase after the first DVE cells have reached the emVE-exVE boundary) (Stower et al. 2023) (Figure 1D’-D’’’). To extract selected VE motion and shape parameters, we used superpixel motion tracking to follow the tissue deformation within each of 32 sub-regions of each embryo (Figure 1E-F, Figure S3, Movie S3-S4) and generated average embryo heat-maps (Figure 1 F’).

We extended our pipeline to find and re-project the corresponding underlying, contoured, basal-surface of the epiblast tissue, as a series of 2D geodesic projections (Figure S1A-B, Movie S1-S2) (Methods). We also generated a novel light-sheet time-lapse data-set where the VE and epiblast of E5.5 embryos were labelled with spectrally orthogonal fluorophores. We did this by crossing the mTmG line (Muzumdar et al. 2007) with a VE-specific TTR-CRE driver line (Kwon and Hadjantonakis 2009) to generate embryos with a GFP-expressing VE and tdTomato-expressing epiblast. The VE and epiblast in these embryos could also be extracted and re-projected separately throughout the time-lapse (n = 3) (Figure S1C, Movie S5-S7). Together, this ∼15 TB time-lapse data-set comprises a total of 194 hours (8 days) of imaging data from 21 embryos (Table S1). Having spatiotemporally aligned, staged and generated an analysis framework, we could now explore the behaviour of each tissue (EPC, VE and epiblast) in the establishment of axial asymmetries.

### Morphological asymmetry in the EPC anticipates A-P axis orientation

The embryo changes in size and shape as it develops. To understand whether there are any asymmetries in the 3D shape of the egg cylinder that might be indicative of A-P axial asymmetries prior to DVE migration, we analysed the 3D volume of our live datasets prior to, and during the course of DVE migration. The morphological asymmetry in the direction of tilt of the EPC has been shown to align with either the anterior or posterior pole of the embryonic axis in fixed samples at E5.5 (Hadjikypri et al. 2024) and E6.5 (Gardner et al. 1992). It is unknown whether this asymmetry precedes DVE migration (Figure 2A) in individual embryos. Having aligned embryos in our dataset along their A-P axis, we could retrospectively determine using our longitudinal live imaged dataset whether the direction of EPC tilt was predictive of A-P axis orientation prior to DVE migration (Figure 2A, Methods). In embryos that showed a morphologically distinct EPC (n = 20/21), we found that the majority (n = 14/20) had an EPC with a pronounced tilt prior to migration (Figure 2B), while the remaining embryos (n = 6/20) showed a straight, proximodistally aligned EPC. In embryos with a pronounced asymmetry in the EPC, we found a bimodal distribution with either a “posterior, left” (n = 8) or “anterior, right” tilt (n = 6) (Figure 2B) so that DVE cells would either migrate in the same direction as the EPC was tilted, or away from it, with an average off-set of 50.79° ± 25.07° across the 14 embryos (Figure 2C). This differed significantly from a random distribution (*χ*^2^ test for deviation from expected uniform probabilities; using 90º sectors, *p*=<0.05, Methods). We also tested whether the tilt of the EPC changed over the course of DVE migration (or whether a straight EPC becomes tilted over time), and found no significant change in the direction of tilt (Figure 2B). This indicates that the asymmetric morphology of the EPC anticipates the orientation of the A-P axis, but is not predictive of the direction of DVE migration.

**Figure 2.**
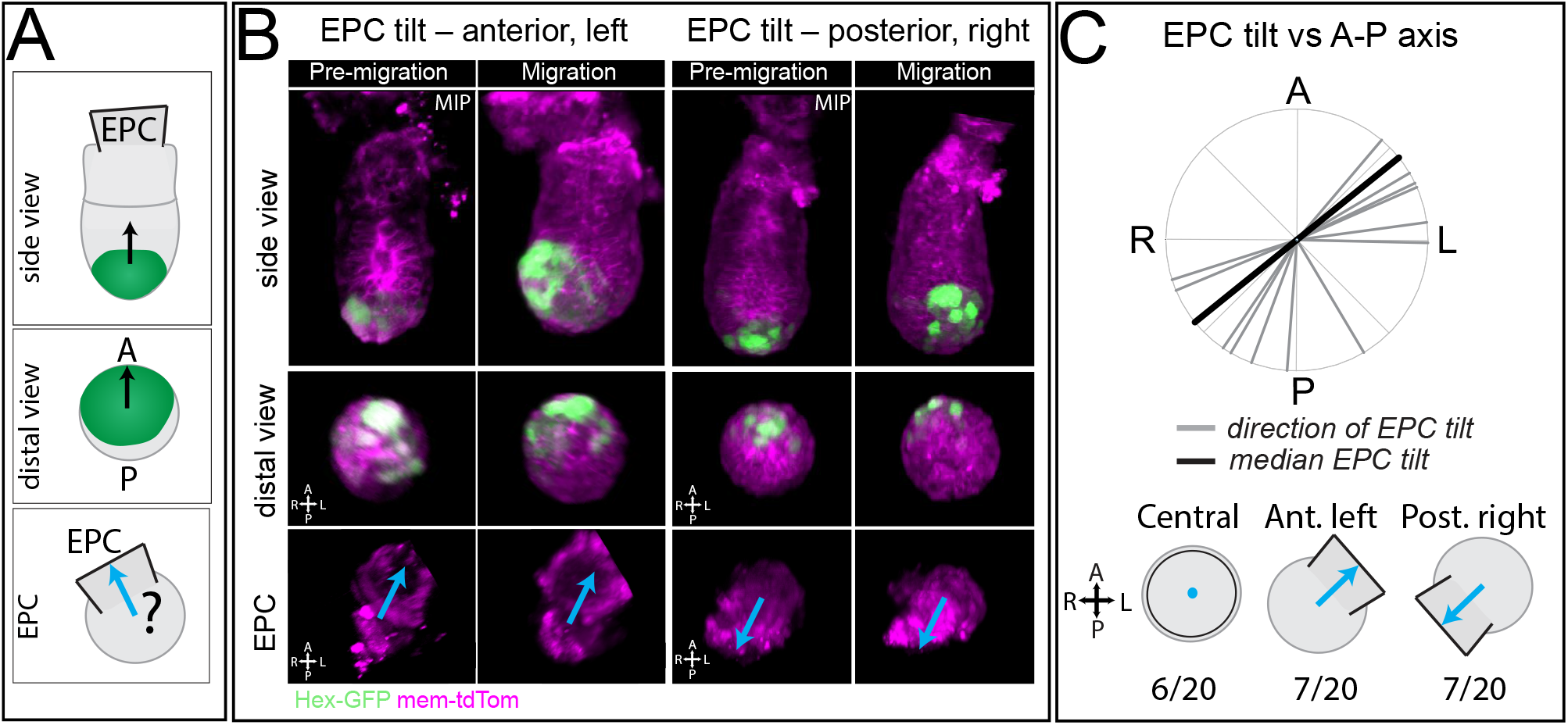
Morphological asymmetry in the ectoplacental cone predicts the position of the A-P axis, but not its polarity. **(A)** Diagram showing EPC in relation to the anterior-posterior (A-P) axis. It is unclear whether the tilt (morphological asymmetry) of the EPC aligns with the A-P axis even prior to DVE migration. **(B)** Example E5.5 embryos imaged by a light-sheet microscope, with examples of EPC-tilted to the anterior-left and posterior right of the A-P axis, showing that DVE cells can migrate away or towards the same side as the EPC. **(C)** Radial plot showing direction of EPC in relation to the A-P axis (black line = median line). Not all embryos showed a clear tilt in their EPC (6/20). In embryos with a morphologically tilted EPC there was an even split between anterior left or posterior right with an average offset of 51°. This differed significantly from a random distribution (*χ*^2^ test for deviation from expected uniform probabilities; using 90º sectors, *p*=<0.05) suggesting that the EPC predicts the axis, but not its polarity (i.e., direction of DVE migration).

### Morphodynamic analysis reveals distinct phases in DVE migration

To understand if there are asymmetries in the behaviour of the VE that precede DVE migration, we used MOSES superpixel-based tracking to measure movement parameters of VE sub-regions of each embryo during each phase of migration (Figure S3-S4, Methods). After spatiotemporal alignment and staging, we found that embryos varied in the duration of DVE migration from the distal tip to the boundary with the exVE (Table S1), with an average time of ∼4 hours (n = 21) similar to previous studies (Srinivas et al. 2004; Takaoka et al. 2011). For analysis of the pre-migration/initiation phase, we used embryos that had a minimum of 1 hour (and average of 3 hours) of time-lapse data prior to migration (n = 14). For analysis of the post-migration phase, we used embryos that had a minimum of 1 hour (and average of 4 hours) of time-lapse data after they had reached the end-point of migration (n = 19) (Table S1). We calculated the rate of: anteriorwards velocity (μm/min, in the anterior direction), total speed (μm/min, in any direction), curl strain fraction (a measure of local rotational motion) and surface area change during each phase (see Methods and Figure S4).

We first computed multi-embryo averaged heat-maps to understand the consensus behaviour across the VE during each phase (Figure 3A, Methods). This confirmed that our tracking was sensitive enough to identify the expected anteriorwards motion of the DVE cell population and the global, bi-rotational pattern of the emVE tissue (Figure 3A, arrows on speed plots, and curl strain plots). We also confirmed that the exVE remained relatively static (Figure 3A, outer rings & Figure S5A), as previously described (Stower et al. 2023; Trichas et al. 2011). To understand how conserved within individual embryos (n = 21) the pattern of local motion behaviour was for each sub-region, we assessed the distribution of motion vectors relative to the ensemble median (Figure S6), using the Rayleigh distribution statistic (*R*) – a measure of angle distribution where *R = 0* is a random distribution, and *R = 1* is uni-directional (Table S2). We find that the average motion pattern is highly conserved across embryos (Figure S6, Table S2). The DVE regions across all embryos (n = 21) have significantly aligned vectors (*R* = 0.92 - 0.97, p = <0.05), higher than surrounding emVE (*R* = 0.49 - 0.80, p = <0.05) (Table S2).

**Figure 3.**
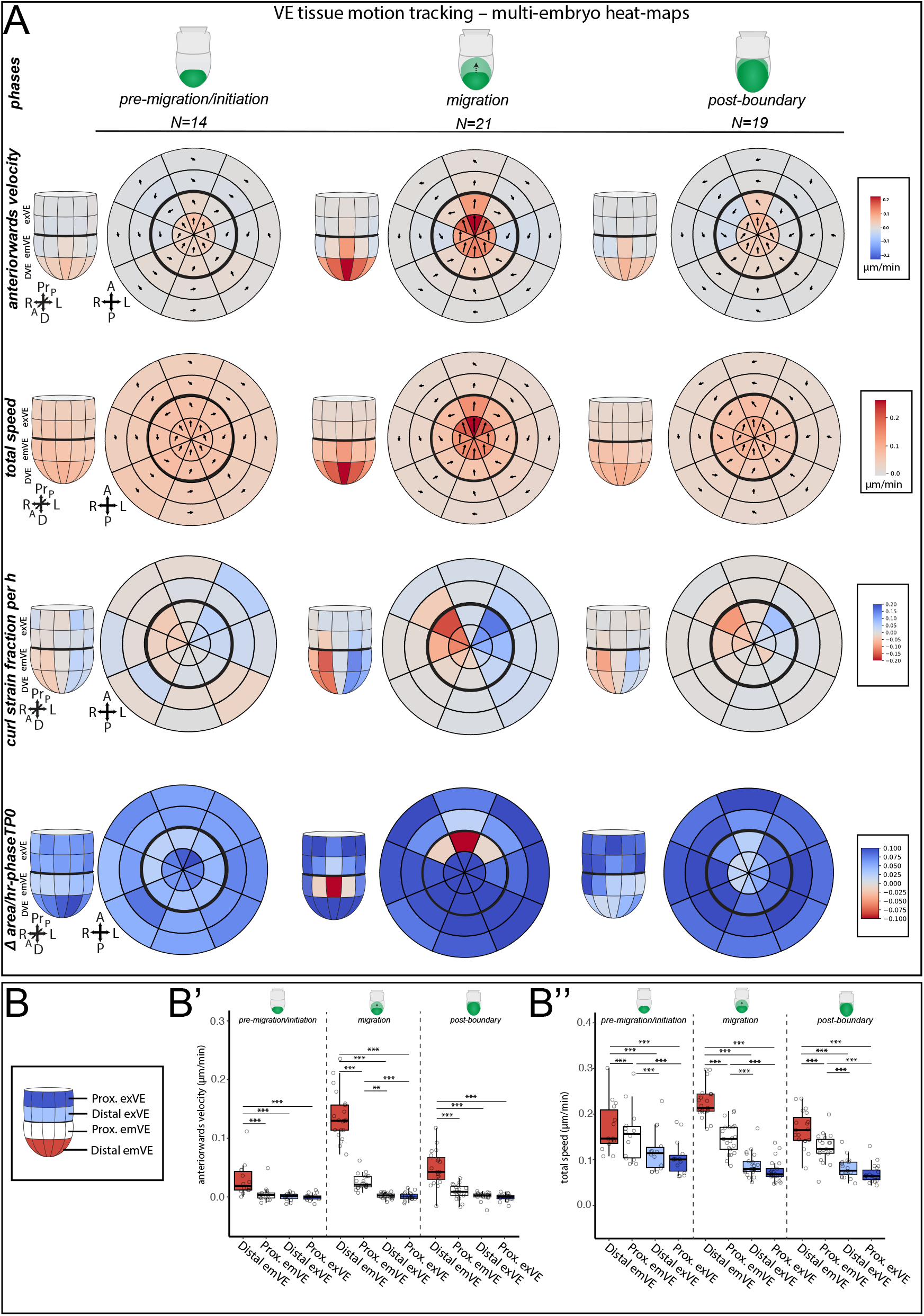
Multi-embryo tissue morphodynamic anaylsis reveals distinct phases in DVE migration. **(A)** Multi-embryo heat-maps of VE tissue motion behaviour showing average behaviour of 32 sub-regions of the DVE of anteriorwards velocity, total speed (in any direction), curl strain fraction per hour, and surface area change. Arrows show median motion vector of superpixels in each region. Arrow length is proportional to the speed value. Central ring is the distal most VE. Black ring denotes the emVE-exVE boundary. Data is converted to 3D coordinate-space for quantification. In motion tracking note the anterior direction of the DVE migration (arrows), and the bi-rotational flow of the surrounding emVE during migration (curl strain fraction). This flow pattern continues into the post-boundary phase where the nascent DVE cells continue to move anteriorly from the distal tip. In surface area change, note that the anterior emVE undergoes a decrease in surface area (orange and red colour) during the migration phase. This decrease in area does not happen during the post-boundary phase. Note that in the ‘pre-migration/initiation’ phase, DVE cell show higher instantaneous anteriorward speed than surrounding regions, but movement is not directionally persistent at this stage. **(B-B’’)** The distal VE regions shows the highest anteriorwards velocity and total speed at migration stages (migration to boundary and post-boundary) (one-way ANOVA, *p*=<0.001, followed by Tukey’s HSD Test for specific comparisons). Interestingly, even in the pre-migration/initiation phase, when the DVE shows low directional persistence (see Figure 1D’), there is still significantly higher motion in the DVE cell population compared to surrounding cells and the exVE (one-way ANOVA, *p*=<0.001, followed by Tukey’s HSD Test for specific comparisons).

Having confirmed that the known patterns of VE motion are captured by this analysis, we next tested whether there was a significant difference in motion or apical surface area in the emVE immediately anterior or posterior to the DVE in the pre-migration/initiation phase that might influence the direction of migration. We found no significant difference in motion behaviour (total speed, anteriorwards velocity) (Figure S5B) or area change (rate of change, fold change) (Figure S5B) between anterior and posterior emVE regions, suggesting that prior to the onset of collective DVE migration, there are no overt *behavioural* asymmetries in motion, or apical surface area change in the emVE that influences DVE migration direction.

We next focused on the behaviour of the DVE cell population. Though the DVE is considered to be largely static prior to the onset of collective migration, in the pre-migration/initiation phase there was motion in the DVE which was significantly higher than surrounding regions, though not of the magnitude of the main migration phase (Figure 3A,B’). This suggests that the DVE is behaviourally distinct in the VE even prior to onset of collective migration. Furthermore, we found that though the DVE continues to move (laterally) once reaching the emVE-exVE boundary as it is replaced by later DVE cells (Takaoka et al. 2011), the motion of this second migratory phase is slower than that of the primary migration phase (Figure 3B-B’’, Figure S5C).

### DVE cell population transitions from disorganised motion to collective, unidirectional ratchet-like migration

MOSES superpixel-based motion tracking analysis revealed that the DVE is behaviourally distinct even prior to the onset of collective migration. However, this analysis measures tissue motion and temporally averages movement within each stage of migration. To understand in greater detail the series of events leading up to, and after, the onset of DVE migration, we analysed individual single-cell movements in our Lifeact-GFP dataset, in which each cell is segmented and its centroid tracked at 5-minute resolution (n = 5 embryos) (Figure 4A-A’) (Stower et al. 2023). We computed the angle histogram of cell velocity with each cell’s contribution weighted by its speed at 10-minute increments during the time period +/-1 hour, relative to the onset of DVE migration (Methods). To quantify the extent of unidirectionality in the histogram, we calculated the Rayleigh distribution statistic (*R)* (Methods). In the pre-migration/initiation phase cells move anteriorwards, but in a less collective manner with wider histogram dispersion (Figure 4A’’), and a relatively low R number (Figure 4B, also see Movie S9 – phase 1). However, at the onset of migration, DVE cell velocity becomes significantly more directionally aligned (Figure 4A’’), evidenced by the Rayleigh distribution statistic increasing by 0.2 after 15 minutes (Figure 4B) (Movie S9 – phase 2). This shows that the onset of DVE migration is not a simple transition from stationary cells to collective unidirectional movement, but rather, of cells becoming more aligned in the direction of migration, resulting in a transition to unidirectional, persistent migration of the DVE population.

**Figure 4.**
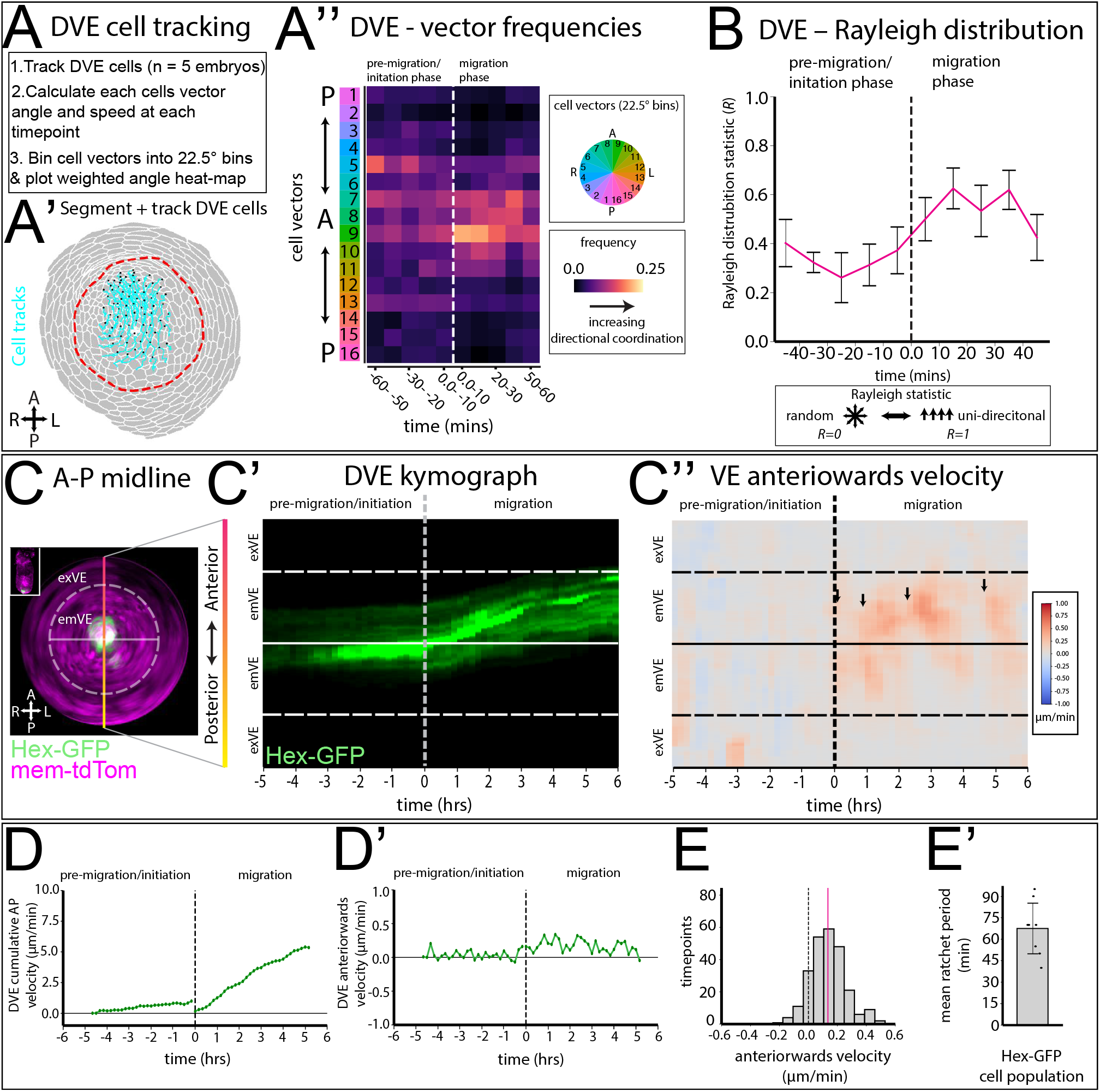
DVE cells collectively migrate in ratchet-like manner. **(A)** Overview of cell movement analysis in cell tracked Lifeact-GFP time-lapse data-set, imaged at a 5-minute interval. Each cell vector angle is assessed then weighted by its velocity. **(A’)** Example embryo showing DVE cell tracks (cyan) on 2D polar geodesic projection. Red dotted line denotes emVE-exVE boundary. **(A’’)** Heat-map of weighted angle histogram of DVE cell velocity where each cell’s contribution is weighted by its speed. Cell vectors are in 22.5° bins for the +/-1 hour from onset of collective migration. Note the increase in coordinated anteriorward motion at the onset of the migration phase (white dashed line). **(B)** Graph showing Rayleigh distribution statistic of DVE cell tracks at pre-migration/initiation and migration phases, which shows the increase in coordination in cell vectors at the onset of migration (0 = random distribution, 1 = uniform distribution). **(C)** 2D geodesic polar projection of an example E5.5 Hex-GFP:mem-tdTomato embryo imaged every 10 minutes for 11 hours. Digitally extracting the A-P midline (coloured vertical line in C) throughout the time-lapse enables a kymograph to be plotted. **(C’)** Kymograph of the Hex-GFP channel along A-P mid-line of the embryo in C, showing the DVE cell population migrating collectively at the onset of the migration phase (vertical dotted line) to reach emVE-exVE boundary (horizontal dotted line). **(C’’)** Kymograph of superpixel motion tracking of A-P mid-line of embryo in C. Plotting anteriorwards velocity shows that the DVE movement is intermittent, undergoing start-stop behaviour (arrows). **(D-D’)** Graphs showing anteriorwards velocity and cumulative velocity (i.e., anteriorwards displacement/ min) of Hex-GFP population of embryo in C. In the pre-migration/initiation phase, where there is a relatively low level of directionally coordinated motion amongst DVE cells (A’’,B), there is little anteriorwards velocity. In the migration phase when directional coordination increased (A’’,B), anteriorwards velocity increases as the DVE migrated collectively (see position of Hex-GFP in D’’. t-5h - t0h). **(E)** Histogram of anteriorwards velocity at all timepoints from 9 Hex-GFP embryos during migration phase. In >90% of time-points Hex-GFP progresses anteriorly, with minimal posteriorwards motion. **(E’)** Plot showing mean time between start-stop events for each Hex-GFP embryo (n = 9); mean ratchet period was 1.13 ± 0.29 hours (mean ± stdev).

To better understand the emergence of this persistent migration, we analysed the Hex-GFP:mem-tdTomato dataset where DVE cells are labelled with GFP (n = 9). We digitally extracted the A-P midline (Figure 4C) and plotted the Hex-GFP channel as a kymograph to visualise the position of the DVE (Figure 4C’) and anteriorward velocity (Figure 4C’’), the latter also being plotted as a line graph (Figure 4D). To assess the effect of local movements with the extent of migration, we also plotted the cumulative AP velocity of the Hex-GFP population (Figure 4D’). These plots show that DVE migration does not proceed with uniform speed, but does so in ‘bursts’ of elevated anteriorwards movement, separated by periods of relative inactivity (Figure 4D-D’). In the pre-migration/initiation phase, there is a relatively low level of directionally coordinated motion among DVE cells (Figure 4A’’, B, Movie S8), resulting in only a small net anteriorwards movement, during which the DVE population remains largely at the embryonic distal tip (Figure 4B’, B’’, Movie S8 – phase 1). In contrast, at the onset of migration, the magnitude of directional coordination increases (Figure 4C’’, D), and the DVE population moves collectively forwards, in ‘bursts’ (Figure 4D’-D’’, Movie S8 – phase 2).

In this manuscript we define ratchet-like behaviour to be movement restricted to one direction with minimal movement in the opposite direction, occurring recurrently. To test whether DVE motion is ratchet-like, we measured anteriorwards velocity from an Anterior-Posterior midline digital-unwrap of the apical VE to the basal epiblast layer (Methods) and plotted a histogram of anteriorwards velocity of all time-points during the migration phase from all embryos (n = 9) (Figure 4E). This revealed that in more than 90% of time-points the DVE shows a net anteriorwards motion, with a mean velocity of 0.2 µm/min. To understand how similar the ratchet-like behaviour is across embryos, we calculated the period between ‘bursts’ for individual embryo by identifying successive peaks in the DVE anteriorwards velocity autocorrelation curves (Methods). This revealed that DVE cells move in bursts with an average recurrence of 1.13 ± 0.29 hours (n = 9) (mean ± stdev) (Figure 4F).

### Vertex model demonstrates that emVE tissue rheology can account for DVE ratchet motion through stress relaxation

To investigate potential mechanisms that could give rise to ratchet-like, start-stop DVE migration, we developed an *in silico* 2D vertex model of the VE tissue (Alt et al. 2017; Fletcher et al. 2014) (Figure 5A, Figure S7A, Methods). Using this model we varied two key parameters: DVE self-propulsion force (*F*), and target shape index of the emVE cells, 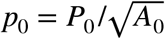 where *P*_0_ and *A*_0_ are the target perimeter and target area of the emVE cells. These parameters control DVE propulsion and rigidity of the emVE cells, respectively (Methods). In alignment with our observations from live *in vivo* data (Stower et al. 2023), DVE cells were given parameters corresponding to a solid-like cell cluster (a small target shape index) with a anteriorwards self-propulsion force (*F*) to generate directional migration (Figure 5A, grey cells), emVE cells were modelled as passive (no self-propulsive force) (Fig 5A, blue cells) and surrounding exVE cells did not evolve in the simulation, thereby defining a boundary (Figure 5A, red cells). Using this model we performed a parameter sweep for ratchet-like movement, measured by a high coefficient of variation in DVE velocity (Methods), running simulations with different magnitudes of DVE propulsive force (*F*) and emVE tissue rigidity (*p*_0_) (Figure 5B, Methods). Our sweep revealed that a region of parameter space supported ratchet-like movement (Figure 5B-B’, Movie S10). We noted that this region of parameter space was also associated with a sudden onset of high level of cumulative T1 transitions (Figure 5B’’). Simulations from this region of parameter space show that the DVE moves as an intact cell cluster (Figure 5A, Movie S10), and confirmed that DVE cells moved with an intermittent, start-stop motion (Figure 5C). We also observed that surrounding cells moved in a polonaise-like flow with counter-rotating vortices (Figure S7B), similar to cell movements *in vivo* (Stower et al. 2023; Takaoka et al. 2011).

**Figure 5.**
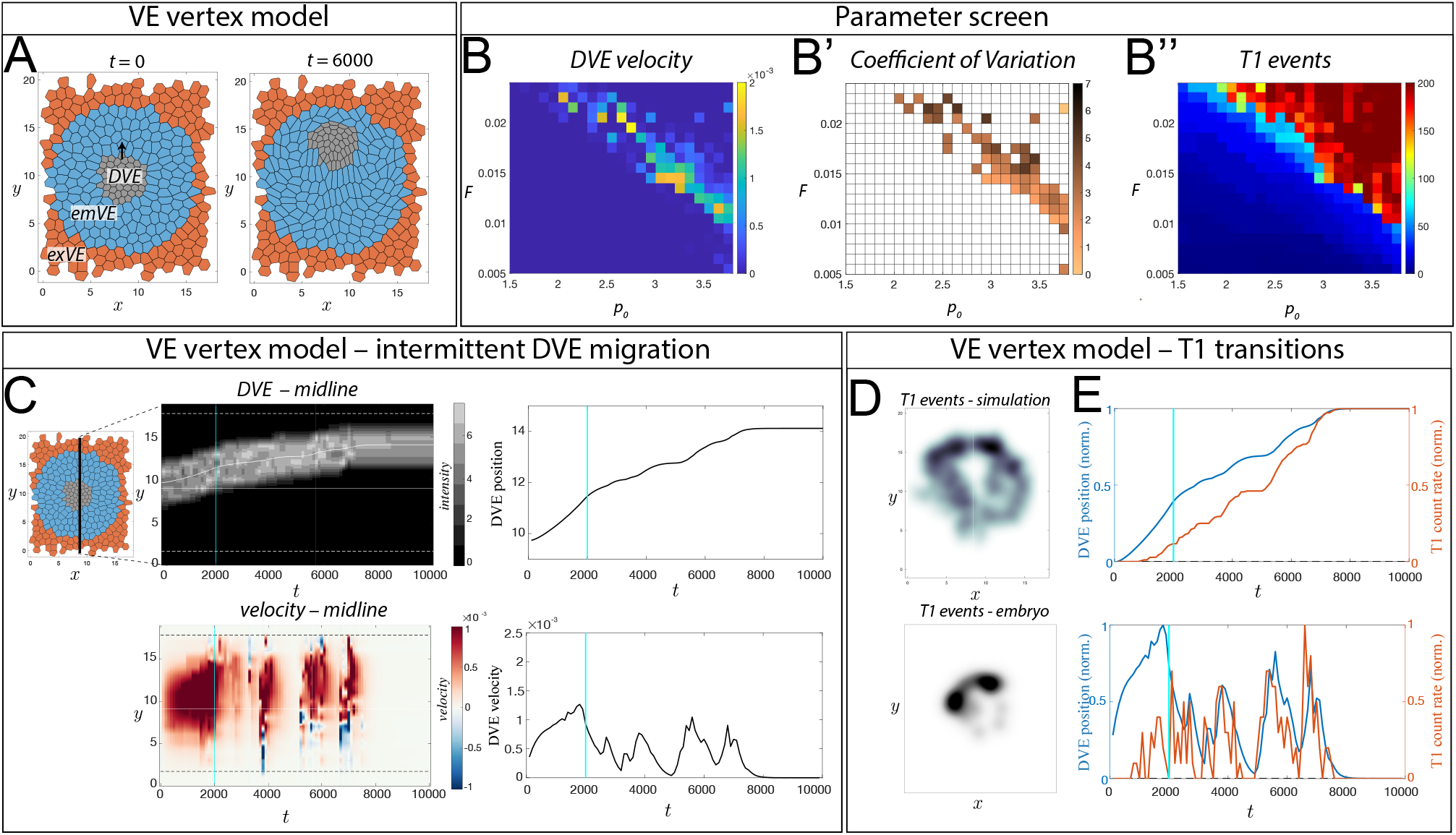
Vertex model shows that DVE start-stop motion is accompanied by T1 transitions in surrounding cells. **(A)** Example of vertex model; DVE cells (grey) were given parameters corresponding to a solid-like cell cluster (a small target shape index) with finite self-propulsion force (*F*). emVE cells (blue) were modelled as passive (no self-propulsive force, *F* = 0) and were less rigid. Surrounding exVE cells (red) do not evolve in the simulation. **(B-B’’)** Parameter space of vertex model formed by varying DVE self-propulsive force (*F)* and rigidity of emVE cells (*p*_0_, target shape index), coloured by: **(B)** maximum DVE velocity, **(B’)** coefficient of variation of DVE velocity, and **(B’’)** cumulative T1 transition events (Methods). We note that only a specific region of parameter space gives rise to intermittent DVE movements. **(C)** Kymograph through the midline of the simulation in A, and line plots showing the position of the DVE centre-of-mass and velocity – confirming that start-stop behaviour occurs in the simulation. **(D)** Density-map of T1 events in example *in silico* simulation (top panel) and *in vivo* Lifeact-GFP embryo (bottom panel), showing that the majority of T1 events occur ahead of the migrating DVE (high density - black, low density - white). **(E)** Temporal analysis of DVE velocity and T1 events in an example *in silico* simulation where *F* = 0.01 and *p*_0_ = 3.6, showing peaks of DVE velocity (blue line) closely aligned with peaks of T1 events (orange line). Also see Movie S10 for an example vertex model simulation.

We next examined the relationship between T1 transitions and start–stop motion in the model, given their established role in stress relaxation during morphogenesis and cell rearrangements (Jain et al. 2023; Krajnc et al. 2018; Zhang 2025). In the solid-phase vertex models, T1 transitions require overcoming an energy barrier, but once neighbour exchange occurs local stresses are reduced, allowing the tissue to transiently flow more readily until stresses build up again (Bi et al. 2015; Krajnc et al. 2018). Analysing the spatial location of T1 events in simulations undergoing intermittent DVE movement, we found that T1 events occurred in the emVE surrounding the DVE, with a higher density in the region ahead of the DVE (Figure 5D - top panel). Interestingly, comparing the timing of T1 events in the model, we found that as DVE movement began there were very few T1 events in surrounding cells (Figure 5E). Subsequently, bursts of T1 transitions appeared to correlate closely with increases in DVE velocity (Figure 5E). Analysis of multiple simulations runs with the same parameter settings and over a range of parameter settings (Figure S7D, Methods) confirmed these two stages are robust in this model; an initial period after DVE migration starts with few T1 transitions, followed by a near instantaneous correlation between T1 events and DVE movement bursts.

In the E5.5 mouse embryo we have previously found that T1 transitions occur almost exclusively in the emVE region ahead of the migrating DVE (Stower et al. 2023). The distribution of T1 transitions in our *in silico* model approximates that found in the embryo (Figure 5D). To understand the relationship between DVE intermittent migration and T1 events *in vivo*, we plotted DVE motion against T1 transitions from our light-sheet imaged data-set (n = 5 embryos). This shows, similar to our *in silico* vertex model, that there is an initial phase of DVE migration with no or very few T1 transitions (Figure S7E). This phase is also followed by DVE start-stop movements associated with bursts of T1 transitions in surrounding emVE cells (Figure S7E). Together our results support a two-stage process of DVE migration; an initial T1 transition-independent stage, where DVE cells can move relatively freely, followed by ratchet-like, intermittent movements coinciding with bursts of T1 transitions in the proximal anterior emVE cells ahead of the DVE cells.

### Anterior epiblast cells show retrograde cellular flow during DVE migration

Having investigated morphological and behaviours asymmetries in the EPC and VE tissue, we next focused on the epiblast, the pseudostratified epithelium underlying the VE (Figure 6A). After alignment with VE cartographic projections (Methods), we performed MOSES superpixel-based tracking of epiblast motion in 2D geodesic projections (Figure 6A-B) to generate a complimentary dataset to that of the VE. This revealed that as DVE cells migrate proximally (Figure 6C,E), the underlying anterior epiblast tissue shows an opposite, distal-wards, planar motion (Figure 6D,E. Figure S8A-B, Movie S11-S12), and the distal-most epiblast undergoes a rotational motion (Figure 6D,E. Figure S8A). This motion behaviour was on average half as fast as that of the DVE (Figure S8C).

**Figure 6.**
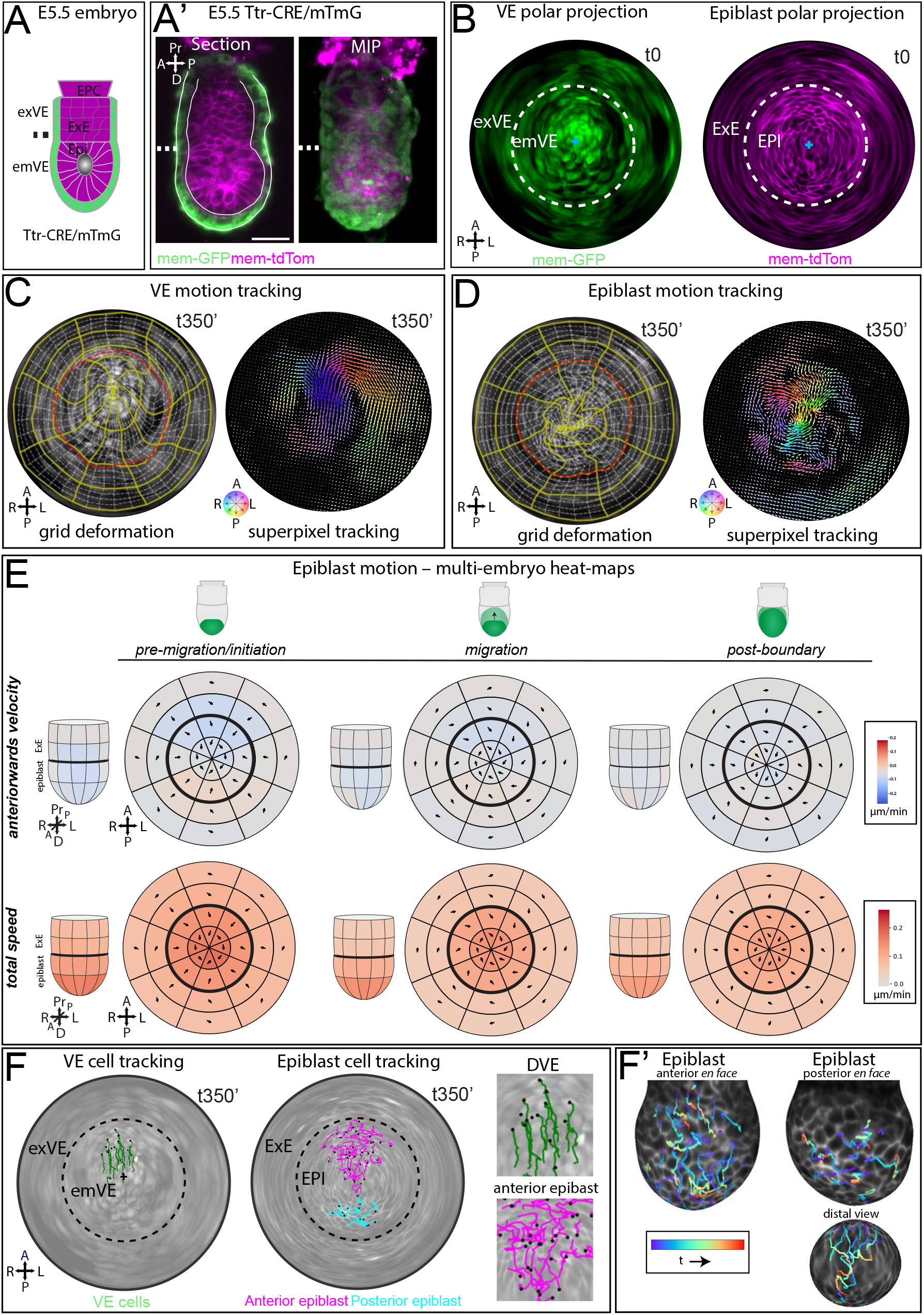
The anterior epiblast shows a distalwards motion during DVE migration. **(A)** Diagram of E5.5 mTmG mouse embryo after Ttr-CRE mediated recombination to label the VE with membrane-GFP and the epiblast and ExE with membrane-tdTomato (magenta). **(A’)** Example of an E5.5 Ttr-CRE/mTmG embryo imaged by light-sheet microscopy showing a single z-plane (left) and a a max intensity projection (right). White dashed line shows emVE-exVE boundary. **(B)** Digital 2D geodesic polar projections of the apical VE (left) and basal surface (right) of the epiblast in the embryo in A’. White dashed line shows emVE-exVE and epiblast-ExE boundary, respectively. **(C)** Superpixel motion tracking of the VE shows the migration of the DVE and the bi-lateral pattern of motion in the VE. **(D)** Superpixel motion tracking of the epiblast during DVE migration shows a posteriorwards flow in the anterior epiblast (yellow-green tracks). Also, see Movies S11-S12 for examples of superpixel tracking. **(E)** Multi-embryo heat-maps of epiblast superpixel tracking during the 3 phases of DVE migration show a flow of motion across the epiblast even during pre-migration/initiation phase. **(F)** Manual tracking of sub-sets of VE (green tracks left panel; black dots denote final position), anterior epiblast (magenta tracks, right panel) and posterior epiblast (cyan tracks, right panel). Black cross denotes distal tip. **(F’)** 3D cell tracking of anterior epiblast and posterior epiblast from embryo in F, here shown with temporally-coloured tracks. Also, see Movie S13 for an example time-lapse with VE and epiblast cell tracking.

The E5.5 epiblast is highly proliferative and cells undergo divisions at their apical aspect (i.e., the surface furthest from the VE). Such division events might be captured as apparent motion behaviour by superpixel tracking in cartographic projections. Therefore, to understand whether our observed motion pattern was due to the planar movement of individual, or groups of, epiblast cells, we additionally hand-tracked cells in the DVE, anterior, and posterior epiblast regions, on the 2D projections of 3 embryos (Figure 6F, Figure S8D-E, Movie S13), and re-projected the tracks to 3D coordinate space (Figure 6F’). This revealed that anterior epiblast cells moved distally during DVE migration (Figure 6F-F’), albeit with lower directional persistence and more convoluted tracks than the DVE (Figure 6F-F’, Movie S13). In contrast, cells in the posterior epiblast show a more limited, random motion (Figure 6F-F’, Figure S8D-E). This indicates that there is a behavioural asymmetry in epiblast cells at these stages.

Given the striking counter-current pattern of motion in the anterior epiblast relative to the DVE, we wanted to confirm this pattern of motion in our data-set. We therefore used extended depth apical-to-basal surface re-projections of the A-P midline of the egg cylinder to extract the motion of both VE and epiblast tissues using superpixel-based tracking (Figure 7A,B). This projection is essentially a mid-sagittal section through the A-P axis that has been computationally ‘straightened’ from a cylinder to a rectangular arrangement allowing the depth of the tissue in this region to be tracked (Figure 7A). We found that the stage-specific, temporal averaged multi-embryo heat maps of this mid-sagittal view show the same DVE motion pattern found in apical cartographic projections and also highlight the retrograde epiblast motion in the anterior epiblast, prior to and during DVE migration (Figure 7C, Movie S14).

**Figure 7.**
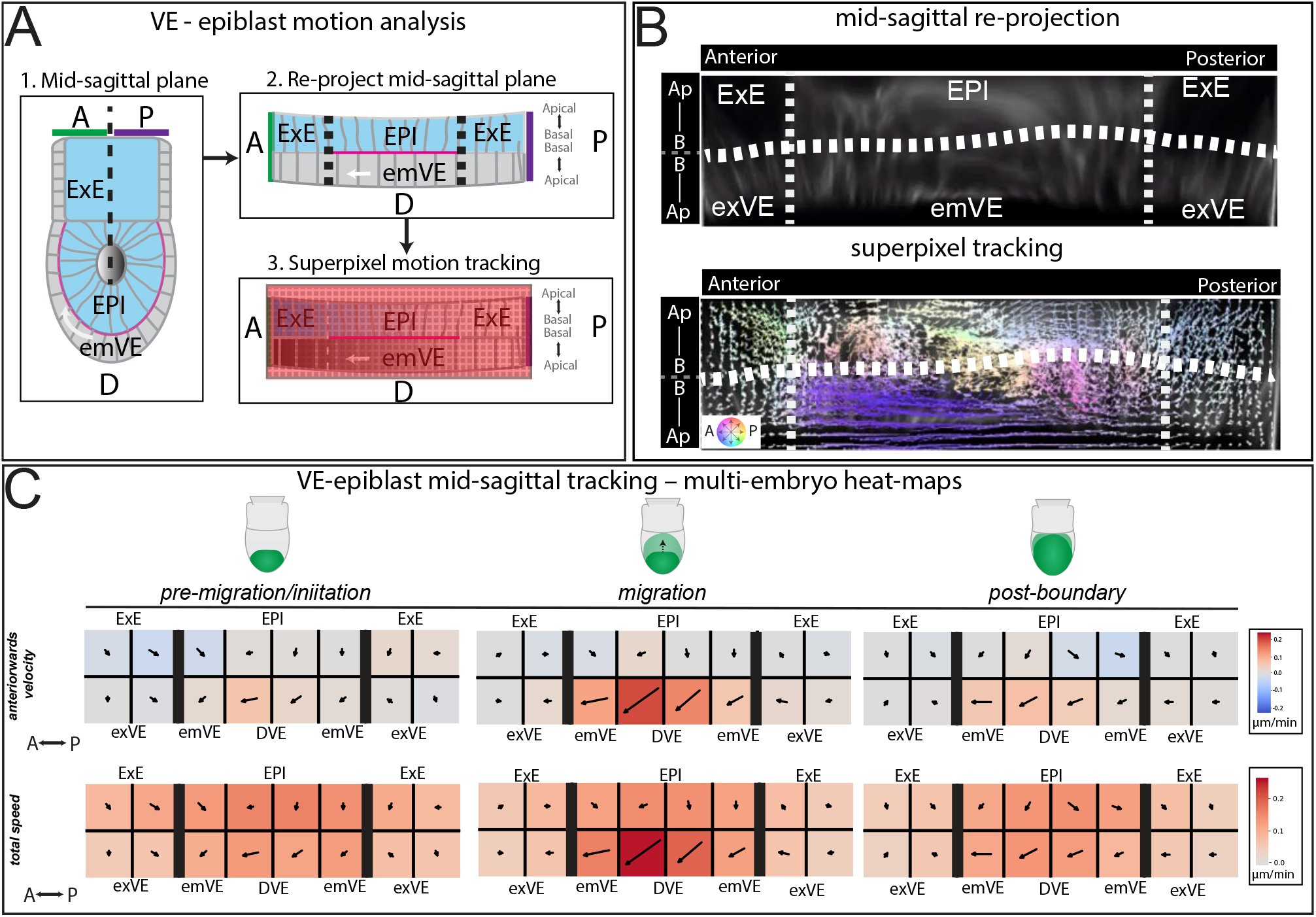
Midline A-P projection analysis shows retrograde motion behaviour. **(A)** Overview of analysis method for digital extraction and linear re-projection of the anterior-posterior mid-line of light-sheet imaged embryos. **(B)** Example linear A-P re-projection of Lifeact-GFP embryo (upper panel) and corresponding superpixel motion-tracking (lower panel) showing the migration of the DVE to the left of the projection (purple tracks), while the epiblast moves to the right (yellow-orange colours). See Movie S14 for an example mid-sagittal re-projection. **(C)** Multi-embryo heat-maps of A-P linear projections showing the average motion behaviour of the epiblast and VE at 3 migration phases in the mid-sagittal tracking analysis.

To maintain tissue integrity, cells adhere to their basement membrane through active remodelling of focal adhesion complexes (FACs), containing Paxillin (Hu et al. 2014; Nayal et al. 2006; Rajah et al. 2019). Given our finding that some epiblast cells can undergo directional planar movements, we asked whether epiblast cells form basal FAC and therefore must actively remodel their adhesion to the basement membrane during this planar motion. Due to the close basal-to-basal apposition of epiblast and VE cells about a shared basement membrane, it is difficult to resolve through immunohistochemistry alone which of the two form FAC (Figure S9A). We therefore made use of a conditional EGFP-Paxillin fusion reporter mouse line (Abe et al. 2011) that allowed EGFP-Paxillin to be expressed in a tissue specific manner in only the VE or epiblast through tissue-specific CRE-drivers (Figure S9B-C). We observed that in all cells of the epiblast (n = 15 embryos) and VE (n = 15 embryos) Paxillin-EGFP localised to the basal cell surface (Figure S9B-C), suggesting that any planar cell movements could involve the dynamic regulation of focal adhesion contacts. Together these data reveal a novel retrograde motion of the epiblast during collective DVE migration and show that epiblast movements must be actively regulated.

## DISCUSSION

The patterning of the A-P axis is a fundamental and essential event in early development. However, it is unclear the extent to which multiple parameters combine to control the asymmetric migration of the DVE in a system that shows natural stochastic variation and is subject to considerable environmental noise. To investigate the morphological and behavioural asymmetries that underlie axis formation, we developed an integrated, multi-scale (cellular, tissue, embryo-level) analysis of motion behaviour in the VE and epiblast in live-imaged embryos. This revealed that multiple levels of inputs may be involved in establishment of axial asymmetries, including morphological and behaviour asymmetries in the epiblast, that exist prior to the onset of DVE migration.

We find a directional planar motion in the anterior opposite to that of the movement of DVE migration – akin to the DVE migrating over a treadmill-like epiblast. Epiblast cells in this region move distally at approximately half the velocity of the overlying DVE cells, while posterior epiblast cells remain relatively static. Previous studies have shown that pharmacological inhibition of Nodal inhibits cell division in the epiblast, but not VE, and results in DVE migration failure (Stuckey et al. 2011). Additionally, embryos chimeric for wild-type and Nodal null cells show a significant accumulation of mutant cells to the anterior epiblast (Lu and Robertson 2004), suggesting that cell movements in the epiblast may not be random. Our study puts the behaviour of the epiblast in the context of the collective migration of the DVE and, together with these previous studies, suggests that the epiblast might not be an entirely passive recipient of patterning cues from the DVE but might participate more directly in the generation of axial asymmetry. This would be consistent with *in vitro* and *in vivo* experiments showing that the epiblast can self-organise. For example, gastruloids that lack VE tissue can break axial asymmetry, but do so with a relatively low efficiency (Harrison et al. 2017). In *Lefty1* and *Cer1* mutants where the DVE does not migrate, a proportion of embryos show axial duplications through the formation of multiple streaks (Perea-Gomez et al. 2002). These data suggest that axial patterning in the embryo might be a multi-step process, with the migration of the DVE reinforcing a ‘noisy’ capacity to self-organise intrinsic to the epiblast.

There are interesting similarities as well as differences between these patterning events in the mouse and birds. Experiments using quail embryos have shown movement within the hypoblast (the equivalent of the mouse VE) that is in the same direction as coordinated movements in the epiblast (Villedieu et al. 2025). In the mouse, as in birds, there is coordinated movement in the epiblast and VE, but the movement in the VE is counter to that seen in the epiblast. In birds, it has been shown that hypoblast cell movements are caused by movement in the epiblast (Villedieu et al. 2025). It will be interesting to see if a similar causal relationship exists in the mouse embryo. In birds, *Nodal* is expressed in the posterior hypoblast and contributes to the induction of the primitive streak within the epiblast (Villedieu et al. 2025). We have previously shown that in the mouse embryo, *Nodal* is expressed within the VE, but in contrast to birds, predominantly in the AVE (Thowfeequ et al. 2024). These differences between the mouse and birds may be due to divergence in patterning mechanisms between mammals and birds, but equally, may also be due to differences in the geometry of the embryo, with rodent embryos being cylindrical and the avian embryo being a flat disk. Comparative studies in other mammals with disk embryos (e.g. rabbit, ruminants, primates, etc), may help to resolve the basis for such differences.

Gene expression asymmetries exist in the VE prior to DVE migration (Yamamoto et al. 2004). Here, leveraging the longitudinal nature of our data set, we assessed whether the direction of tilt of the ectoplacental cone, prior to DVE migration, correlated with the future A-P axis. We found that the tilt of the EPC correlated with the orientation of the A-P axis, but not its polarity. This extends previous findings in fixed mouse embryos that suggested that the tilt of the EPC aligned with the orientation, but not the polarity, of the A-P axis in E6.5 (Gardner et al. 1992) and 5.5 (Hadjikypri et al. 2024) embryos. EPC tilt has been suggested to originate from the morphological asymmetry of the pre-implantation blastocyst. This, together with the gene-expression asymmetries present in the primitive endoderm of the blastocyst (Takaoka et al. 2006; Takaoka et al. 2011), supports a model where A-P axial asymmetry exists prior to DVE migration, but the DVE, through its migration, reinforces and amplifies these early asymmetries, ensuring the reliable establishment of a ‘definitive’ A-P asymmetry by positioning the primitive streak.

As the DVE secretes inhibitors of mesodermal fate, our finding that planar epiblast motion occurs at pre-gastrulation stages may result in a greater proportion of epiblast cells exposed to secreted DVE signals. Such an interaction between patterning and cell movements amplifying the effect of signalling has previously been proposed in the node and tail-bud (Fulton et al. 2022). In this context it would would act to increase the range of DVE signalling, but could also act as a timing control system, preventing the precocious formation of mesoderm, and even providing enough time to generate sufficient epiblast cells to then start differentiating during gastrulation.

Our detailed examination of how DVE movements lead to the cells becoming asymmetrically positioned in the embryo revealed unexpected behavioural complexity. We find that prior to the onset of anterior migration, DVE cells show relatively uncoordinated movements before abruptly transitioning to coordinated unidirectional movement, which marks the onset of migration. This indicates that cells within the DVE population are already motile prior to the onset of anterior/proximal migration, and that this motility becomes rapidly channeled to become directional at the onset of migration. What triggers this transition from random to directional migration remains unknown.

Furthermore, we find that once DVE cells start moving collectively they do so in a ratchet-like manner, with recurrent bursts of elevated anteriorwards movement. An i*n silico* vertex model shows that start-stop behaviour is correlated with T1 transitions in surrounding emVE cells and we found a similar correlation in the embryo. Interestingly, both *in silico* and *in vivo* data show that initially, DVE movements occur without T1 transitions in the surrounding cells. Together, these observations suggest that intermittent DVE migration during the main migration-to-boundary phase emerges from an interplay between active DVE motility and the rheological resistance of the surrounding emVE tissue. Resistance from the emVE cells leads to periods of slowed motion, followed by local stress relaxation through cell rearrangements, allowing renewed forward movement until resistance again accumulates. Qualitatively similar intermittent dynamics are also observed during sedimentation of spheres of different densities in complex fluid with microstructures that exhibit yield stress rheology (Sgreva et al. 2020; Sgreva and Davaille 2024). This system is analogous to the DVE migration whereby the DVE cluster is a solid sphere, the unidirectional DVE migration is the gravitational force on the solid sphere, and the yield-stress-like rheology of the emVE cells through which the DVE cluster migrates, mimics the complex fluid through which the solid sphere sediments (Amiri et al. 2023; Nguyen et al. 2025).

We also note that, in the embryo, substantially fewer T1 transitions are observed than predicted by the vertex model, and these are largely confined to the region ahead of the DVE. This discrepancy may reflect additional modes of stress relaxation available *in vivo*, as emVE cells can undergo pronounced shape deformations (Stower et al. 2023; Trichas et al. 2011) and cell division events (Stower et al. 2023). Consequently, in the embryo, emVE cells likely employ multiple mechanisms to accommodate the stresses generated by active DVE migration. Looking ahead, an important challenge will be to uncover the molecular and cellular bases of these distinct phases of DVE migration, and to understand in more detail how the behaviours of surrounding cells are dynamically coordinated to support robust DVE movement within the VE epithelial sheet.

## Supporting information

animated abstract

## ACKNOWLEDGEMENTS

We thank Dr Satish Arcot Jayaram for help with mouse breeding, Dr Tristan Rodriguez for the Hex-GFP line, Dr Roland Wedlich-Söldner for the Lifeact-GFP line, Dr Liqun Luo for the ROSA26^mTmG^ line, Dr Toshihiro Fujimori and Riken Center for Developmental Biology for the ROSA26^EGFP-Paxillin^ line, and Dr Anna-Katerina Hadjantonakis for the Ttr-CRE line. This work was funded by BBSRC Research Grant BB/J00989X/1, BBSRC ALERT13 Award BB/L014750/1, Wellcome Strategic Award 108438/Z/15/Z, Senior Investigator Award 103788/Z/14/Z and Discovery Award 227926/Z/23/Z to SS. FZ and XL were funded by the Ludwig Institute for Cancer Research. HH was funded by Wellcome Trust Chromosome and Developmental Biology Doctoral Training Programme 109100/Z/15/Z. JR acknowledges financial support from the Slovenian Research and Innovation Agency (development funding pillar RSF-0106 and research core funding P1-0055). RV acknowledges the support of the Leverhulme Trust, Grant No. LIP-2020-014. RV and JY acknowledge the support of ERC Advanced Grant ActBio, funded as UKRI Frontier Research Grant EP/ Y033981/1). We thank the Micron Advanced Bioimaging Facility, supported by Wellcome Strategic Awards 091911/B/10/Z and 107457/Z/15/Z. We thank Pathology Services Building and Biomedical Services staff for excellent animal support.

## AUTHOR CONTRIBUTIONS

Conceptualisation: SS, FYZ, MJS, RV, JR, JMY

Light-sheet live imaging: MJS

Superpixel-based tracking software development: FYZ

Superpixel-based tracking analysis: FYZ, MJS

Vertex model development and analysis: RV, JR, JMY

EPC angle analysis: HH, MJS

Immunofluorescence and fixed imaging: MJS

Data analysis: FYZ, HH

Mouse husbandry: JG

Writing: MJS, FYZ, RV, JR, JMY, SS

Resources and funding: SS, JMY, ZL, JR

## SUPPLEMENTAL MOVIE LEGENDS

**Movie S1. E5.5 Hex-GFP:membrane-tdTomato light-sheet time-lapse and geodesic tissue projections**. Example E5.5 mouse embryo expressing Hex-GFP in migratory DVE cells and membrane-td-Tomato, ubiquitously. Full volume z-stacks from 2 view angles (0° and 180°) were acquired at a 10 minute interval in a ZIESS Z.1 light-sheet then spatiotemporally registered and fused. For geodesic projections the outer surface of the VE epithelium, and basal contour of the epiblast tissue were computationally extracted and re-projected throughout the time-lapse. Embryos were aligned along their A-P axis with anterior to the top for consistent analysis across samples.

**Movie S2. E5.5 Lifeact-GFP light-sheet time-lapse and geodesic tissue projections**. Example E5.5 mouse embryo expressing Lifeact-GFP. Full volume z-stacks from 2 view angles (0° and 180°) were acquired at a 5 minute interval in a ZIESS Z.1 light-sheet then spatiotemporally registered and fused. For geodesic projections the surface of the VE epithelium, and basal contour of the epiblast tissue were computationally extracted and re-projected throughout the time-lapse. Embryos were aligned along their A-P axis with anterior to the top for consistent analysis across samples.

**Movie S3. 5.5 Hex-GFP:membrane-TdTomato VE polar projection time-lapse, DVE classifier and superpixel motion tracking**. Example E5.5 Hex-GFP:membrane-tdTomato embryo time-lapse geodesic polar projection of the VE epithelium. Superpixel motion tracking of Hex-GFP+ve DVE cell population across embryos enabled a DVE motion classifier to be trained to identify DVE motion phases. At the start of movies a 32-sector a grid was seeded to equidistantly partition the embryonic and extra-embryonic regions. The grid was then deformed by superpixel motion tracking enabling the local behaviour of the tissue to be quantified during each phase of migration.

**Movie S4. 5.5 Lifeact-GFP VE polar projection time-lapse with DVE classifier and superpixel motion tracking**. Example E5.5 Lifeact-GFP embryo time-lapse geodesic polar projection of the VE tissue. DVE motion classifier pre-trained on Hex-GFP data-set was applied to Lifeact-GFP time-lapse data to identify 3 phases of migration. VE superpixel motion tracking was used to deform a 32-sector grid for quantification of local tissue behaviours across the tissue.

**Movie S5. Example timepoint of live imaged E5.5 dual-colour embryo**. Dual colour embryos were generated through a Ttr-CRE:mTmG cross so that the VE epithelium is labelled with membrane-targeted eGFP through a CRE-mediated recombination event, while the epiblast and extra-embryonic ectoderm tissues continue to express membrane-targeted td-Tomato. Movie shows a Z-stack fly-through and a 3D-opacity rotation of half the image volume.

**Movie S6. E5.5 Dual-colour embryo live time-lapse and geodesic tissue projections**. Example E5.5 mouse embryo expressing membrane-GFP in VE cells and membrane-td-Tomato in the epiblast and ExE tissues. Full volume z-stacks from 2 view angles (0° and 180°) were acquired at a 10 minute interval with a ZIESS Z.1 light-sheet microscope then spatiotemporally registered and fused. For geodesic projections the surface of the GFP+ve VE epithelium, and basal contour of the tdTomato+ve epiblast and ExE tissues were computationally extracted and re-projected throughout the time-lapse.

**Movie S7. E5.5 membrane-GFP VE polar projection time-lapse with DVE classifier and superpixel motion tracking**. Example E5.5 Ttr-CRE-mTmG embryo time-lapse geodesic polar projection showing the membrane-GFP+ve VE tissue only. DVE motion classifier pre-trained on Hex-GFP data-set was applied to the time-lapse dataset to identify phases of migration. VE superpixel motion tracking was used to deform a 32-sector grid for quantification of local tissue behaviours across the tissue.

**Movie S8. Onset of DVE collective migration in Hex-GFP:tdTomato embryos**. Example E5.5 Hex-GFP:mem-TdTomato embryo imaged using a ZIESS Z.1 light sheet microscope prior to, and during the onset of DVE migration with the VE tissue polar projected. In Phase 1: pre-migration/initiation phase, there are cell movements but a low level of directionally coordinated motion among the DVE. In Phase 2: the magnitude of directional coordination increases and the population moves collectively anteriorly. Lower panels show a magnified version of the polar projections showing the tdTomato and Hex-GFP channels.

**Movie S9. Onset of persistent DVE directional migration in LifeactGFP embryo with cell tracking**. Example E5.5 Lifeact-GFP embryo imaged using a ZIESS Z.1 light sheet microscope prior to, and during the onset of DVE migration with the VE tissue polar projected and VE cells segmented and tracked. In Phase 1: pre-migration/initiation phase, there are cell movements but a low level of directionally coordinated motion among the DVE cells. In Phase 2: the magnitude of directional coordination increases and the population moves collectively anteriorly. Lower panels show a magnified version of the upper geodesic polar projections.

**Movie S10. Vertex model of ratchet-like DVE migration**. Top left: Simulation of the 2D vertex model showing the DVE cluster (grey) migrating through surrounding emVE cells, with colour indicating local cell shape index (blue indicates low values whereas yellow indicates high values). Top right: Distribution of T1 events (blue region) during DVE migration. Black line on each cell denotes the axis of elongation. Bottom left: Time series of DVE centre-of-mass velocity showing intermittent, burst-like (ratchet) motion. Bottom right: Cumulative number of T1 transitions over time.

**Movie S11. E5.5 Dual-colour embryo live time-lapse and geodesic tissue projections with VE and epiblast superpixel motion tracking**. Example E5.5 VE-labelled membrane-GFP and epiblast-labelled membrane-tdTomato geodesic polar projected lightsheet time-lapse showing the outer surface of the VE epithelium and epiblast side-by-side. Using superpixel motion tracking to deform an 32-sector equidistant grid, local and global tissue behaviours in the VE and epiblast can be directly compared in each embryo during each phase of DVE migration. While the DVE moves anteriorly in the VE epithelium, the flow of the epiblast underneath moves in an opposite posterior direction.

**Movie S12. E5.5 Hex-GFP:mem-tdTomato embryo live time-lapse and geodesic tissue projections with VE and epiblast superpixel motion tracking**. Example E5.5 Hex-GFP:mem-tdTomato geodesic polar projected light-sheet time-lapse showing the outer surface of the VE epithelium and epiblast side-by-side. Using superpixel motion tracking to deform an 32-sector equidistant grid, local and global tissue behaviours in the VE and epiblast can be directly compared in each embryo during each phase of DVE migration. While the DVE moves anteriorly in the VE epithelium, the flow of the epiblast underneath moves in an opposite posterior direction.

**Movie S13. VE and epiblast cell tracking during DVE migration**. Example E5.5 Ttr-CRE-mTmG embryo time-lapse geodesic polar projection showing the VE labelled with membrane-GFP and the epiblast labelled with membrane-tdTomato. Cells in the DVE (green tracks), anterior epiblast (magenta tracks) and poster epiblast (cyan tracks) were hand-tracked. As the DVE cells migrate in the VE epithelium (green tracks) the anterior epiblast underneath moves posteriorward (magenta tracks), while posterior epiblast remain localised (cyan tracks).

**Movie S14. VE and epiblast show counter flow during DVE migration**. Example E5.5 Lifeact-GFP embryo time-lapse showing an extended depth, apical-to-basal, surface re-projection of the A-P midline of the embryo. The projection is a mid-sagittal section through the A-P axis that has been computationally ‘straightened’ from a cylinder to a rectangular arrangement with anterior to the left, allowing the depth of the tissue in this region to be tracked (See Figure 7A for diagram). As DVE cells move anteriorly (left in the projection), epiblast shows an opposite, posteriorwards direction (right in the projection).

## METHODS

### Ethics statement

All experiments were approved by Oxford University Departmental Animal Welfare and Ethical Review Board (AWERB) and were performed under UK Home Office Licence Project 30/3420 Protocol 03 Breeding and Maintenance of genetically altered animals (Mild), and UK Home Office Licence Project PP7051082 Protocol 05 Breeding and Maintenance of GA animals (Mild).

### Mouse Strains, Husbandry, and Embryo Collection

Lifeact-GFP (n = 9) and Hex-GFP:membrane-tdTomato (n = 9) light-sheet imaging data-sets have been previously described (Stower et al. 2023). For additional live imaging experiments to generate embryos where the VE and epiblast were labelled by membrane-GFP and membrane-tdTomato, respectively. Ttr-Cre (Kwon and Hadjantonakis 2009) were crossed with ROSA26^mTmG^ (Muzumdar et al. 2007). In this cross membrane-tdTomato is expressed in all cells, until Cre-mediated excision of the membrane-tdTomato and associated stop codon, enables downstream expression of membrane-targeted GFP construct in the VE. For Paxillin immunohistochemistry Hex-GFP stud males were crossed with CD1 females (Charles River) and dissected at E5.5. ROSA26^EGFP-Paxillin^ (Abe et al. 2011) were crossed with TTR-Cre (Kwon and Hadjantonakis 2009) or Sox2-Cre (Hayashi et al. 2002) mice to drive expression of the EGFP-Paxillin reporter in the VE and epiblast, respectively. Mice were maintained on a 12 hour light, 12 hour dark cycle and noon on the day of finding a vaginal plug was designated 0.5 days *post coitum*. Embryonic day 5.5 (E5.5) embryos were dissected in M2 medium (Sigma) with fine forceps and tungsten needles and transferred into pre-heated culture medium (as per (Trichas et al. 2011)) supplemented with antibiotics, and placed in an incubator at 37°C, 5% CO_2_ prior to imaging, or fixed in 4% PFA for immunohistochemistry.

### Immunofluorescence

Embryos were fixed in 4% PFA at room temperature for 20 min, washed at room temperature three times for 5 min each in 0.1% Triton-X100 in PBS; incubated in 0.25% Triton-X100 in PBS for 25 min; washed three times in 0.1% Tween-20 in 1xPBS; blocked with 2.5% donkey serum, 2.5% goat serum, and 3% Bovine Serum Albumin (BSA) in 0.1% Triton-X100 in PBS overnight; then incubated overnight at 4°C in primary antibodies diluted 1:100 in blocking solution. Embryos were washed three times in 0.1% Tween-20 in PBS (PBT) for 5 min each, with a final additional wash for 15 min; incubated overnight at 4°C with appropriate secondary antibody 1:100 in 0.1% PBT; embryos were incubated with phalloidin at 1nM concentration in PBT overnight at 4°C, washed four times for 5 min in PBT at room temperature; and mounted with Vectashield mounting media containing 4′,6-diamidino-2-phenylindole (DAPI) (Vector Labs H-1200).

### Antibodies and phalloidin staining

Primary antibodies used were rabbit anti-Paxillin (Millipore, 2234097). Secondary antibody used were 1:100 Alex-Fluor (AF)-555 donkey anti-rabbit (Invitrogen, A31570). For F-actin staining phalloidin-atto 647N (Sigma, 65906) was used at a 1nM final concentration in 0.1% Tween20 in 1 x PBS.

### Confocal microscopy of fixed embryos

Fixed embryos were imaged on a ZEISS LSM 880 confocal microscope using a 40x oil (1.36NA) objective. Z-stacks of embryos were acquired at 1 µm interval using non-saturating parameters. Figures were prepared with Adobe Photoshop and Adobe Illustrator (Adobe Inc.).

### Light-sheet time-lapse imaging of Ttr-Cre:mem-tdTomato embryos

E5.5 embryos were placed within a 150 µm-diameter lumen of an agarose cylinder and mounted in a ZEISS Z.1 light-sheet microscope, as previously described (Stower et al. 2023). The width of the lumen in the agarose cylinder was selected as it is wide enough to not compress E5.5 embryos and enables them to remain upright within the imaging chamber. Embryos were imaged using a plan-apochromat 63X/1.0 NA water immersion lens. Full volume z-stacks 1920 x 1920 pixels, 16-bit resolution of each embryo were obtained at a 2 µm interval, recorded sequentially from 2 imaging angles (0° and 180°), with a 135 µW ± 228 nW 488 nm (Coherent) and a 106.9 µW ± 248 nW 561 nm laser (Coherent) 106.9 µW ± 248 nW for GFP and td-Tomato, respectively. eGFP and tdTomato channels were obtained in parallel using a 561 LP secondary beam splitter. Each z-plane was illuminated sequentially with right and left illumination pivot scans oscillating at 23 kHz. The laser light-sheet was focused through a pair of 10X/0.2 NA lenses and the paired lateral illuminations were fused using ZEN Black (ZEISS) “online dual-fusion” setting. A pair of 2-channel z-stack volumes were acquired every 10 minutes.

### Quantitative processing of raw embryo videos

Each embryo time-lapse was processing using our previously established workflow described in detail in Stower et al., (Stower et al. 2023). For each video we i) construct a spatiotemporally consistent 3D coordinate system, ii) unwrap into 2D and spatially align the VE and proximal underlying epiblast surface, and iii) extract tissue motion, stage DVE migration and align migration relative to anterior-posterior axis.

### Construction of spatially and temporally consistent 3D embryo coordinates

Embryo time-lapse videos were processed using the STrEAMS framework as described in Stower et al. (Stower et al. 2023). Briefly, the raw two-angle, z-stack video sequence acquired using a ZEISS Z.1 light-sheet microscope was preprocessed as follows. Images were interpolated to make voxels spatially isotropic, pixel intensities were then converted from 16-bit to 8-bit and volumetrically downsized by a factor of 2. The preprocessed dataset was then spatiotemporally registered; first aligning acquisition angle 1 temporally using the first time-point as the reference image for all time-points, aligning acquisition angle 2 to the registered angle 1 sequence in every time-point, and fusing both angles. Lastly, non-rigid registration is used to match the outer embryo shape over time to the shape at a reference time-point half-way through the time-lapse. For the two-colour acquisitions (eGFP and td-Tomato), we use STrEAMs with the epiblast expressing membrane-targeted tdTomato channel which stains the whole embryo though with weaker fluorescence in the VE and apply the learnt geometric transformations to the second VE expressing membrane-targeted GFP and Hex-GFP channels respectively.

### VE and epiblast surface unwrapping and alignment

The VE layer and its underlying adjacent epiblast was unwrapped from 3D-to-2D using the spline-based method of Stower et al. (Stower et al. 2023) to visualise the cell dynamics in Cartesian, as well as, polar geodesic projections for subsequent quantitative analysis. Due to the use of spatial-temporal registration to construct a consistent 3D coordinate framework over time, the unwrapping transform needs to only be generated for the initial timepoint, t1, then iterated across all subsequent time-points in the time-lapse.

### Specifying the VE and epiblast surface-of-interest

The 3D coordinates of the apical surface of the VE and basal surface of the epiblast to project were manually annotated and specified as binary volumes using the initial timepoint, t1. We found annotating the epiblast surface sub-basally (i.e, just off-set from the epiblast-VE interface) enabled the visualisation of individual epiblast cells in the unwrapped projections.

### Spatial alignment of VE and epiblast surface 2D projections

The shape of the VE and epiblast surfaces were generally not the same but similar in convexity. We thus align the two surfaces by rescaling the unwrapped epiblast projections to be the same pixel size as the unwrapped VE projections. This is conceptually equivalent to doing a nearest neighbour shape alignment without overlap.

### Superpixel-based tissue motion extraction, embryo staging and anterior-posterior alignment

#### Tissue motion extraction

We applied motion sensing superpixels with DeepFlow computed dense optical flow. Superpixel motion tracks at 1000 superpixels (larger region-of-interest) and 5000 superpixels (smaller region-of-interest) on both 2D Cartesian and polar geodesic projection time-lapses of each embryo were extracted.

#### DVE Migration staging

A support vector machine classifier pre-trained on Hex-GFP:mem-tdTomato embryos (Stower et al. 2023) was applied to the tracks from 1000 superpixels to identify DVE-associated superpixel tracks. This classification was then transferred to the tracks from 5000 superpixels and used to compute a consensus DVE migration stage using Cartesian and polar geodesic 2D projections based on directional persistence as previously described (Stower et al. 2023).

#### Determining consensus DVE migration angle

The mean curvilinear 3D velocity of the most directionally persistent subset of the DVE-associated superpixel tracks from 5000 superpixels was used to estimate the DVE migration angle (Stower et al. 2023). The mean of the estimated DVE angle from Cartesian and polar projections defined the consensus migration angle.

For two-channel videos, the epiblast expressing membrane-targeted tdTomato channel associated tracks which simultaneously captures VE and epiblast motion comparable to Lifeact-GFP was used for migration staging and DVE migration angle determination.

### Measurement of EPC angle

Measurement of ectoplacental cone (EPC) angle with respect to the embryonic A-P axis was carried out as previously described (Gardner et al. 1992), but here using the direction of DVE migration to define the A-P axis. Light-sheet time-lapses of E5.5 embryos were spatiotemporally registered, the VE surface extracted, and reprojected as a series of geodesic polar projections. Motion sensing superpixel tracking enabled the directionally persistent DVE-classified tracks to compute mean direction (DVE migration direction), and this was applied to the 3D data. Having aligned light-sheet data by DVE migration (A-P axis), 3D-rendered proximal views of each embryo enabled the offset (0° - 360°; 0° = aligned with anterior, 180° = aligned with posterior) alignment of the EPC from this axis (EPC angle) to be measured prior to DVE migration. The EPC angle of each embryo was also assessed post-migration.

### Quantitative multi-embryo MOSES tissue motion analysis

Individual embryos are heterogeneous in size, shape and rate of growth. The DVE also migrates for different durations. To enable consistent statistical inter-comparison of DVE migration across embryos, we manually outlined the embryonic VE – extra-embryonic VE (em-ex) boundary as a consistent landmark and partitioned each embryo into spatial sectors relative to DVE migration angle and the em-ex boundary (Stower et al. 2023). Crucially these spatial sectors are consistently labelled such that sector 1 always references the distal-most sector that symmetrically overlaps the DVE migration angle. Sector IDs then increase clockwise by increasing radial angle, and by increasing geodesic distance from the distal tip (Figure S3A). This enables statistical tests of sub-regions to be made across embryos in a consistent manner.

### Average embryo superpixel motion heat maps

The average embryo motion heat maps (Figure 3) record the mean statistic of a quantitative measure across 32 spatial sectors for a migration stage. The sectors are constructed in a consistent manner (see below) to refer to the same relative spatial region in every embryo. This allows ensemble averaging of this map on a per sector basis across multiple embryos to construct the average behaviour and perform statistical significance tests. In this paper, we visualise the values of the map using the ‘coolwarm’ colour scheme for signed measures that can have both positive and negative values such as anteriorwards velocity and curl strain, and the ‘reds’ colour scheme for unsigned measures such as total speed which take only positive values.

### *Partitioning the VE and epiblast into sector region-of-interests (ROIs)* for statistical analysis

Each embryo was partitioned in the polar geodesic projection into 32 sector regions-of-interest; 8 radial angle bins that equally partition the radial space of radians, and 4 geodesic distance bins, which partitions the geodesic distance, from the central point (distal tip of the embryo). The first two distance bins equidistantly partitions the distance between the central point (distal tip of the embryo) and the em-ex boundary. The last two distance bins equidistantly partitions the distance between the emVE-exVE boundary and an outer concentric ring of distance, 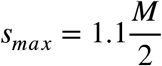. Where ***M*** is the image dimensions of polar unwrap (M x M pixels). This spatial partitioning for each embryo was performed using the first frame of the migration phase.

The same 32 sector partitioning for the VE is used for the underlying epiblast, after spatially aligning the VE and epiblast unwrapped projections (see above).

### Tissue motion analysis

To characterise the net tissue-level motion we used Motion sensing superpixels to seed an initial regular grid of 5000 superpixels and tracked how they were moved by the motion of VE and epiblast cells during each migration stage. The degree of change over time captures the cumulative effect of all cellular events including cell movements, cell shape changes, cell division events and cell mixing. For each superpixel track three measures of average speed were analysed; total curvilinear, mean curvilinear and anteriorwards velocity (see Figure S4). For the computation of all these measurements, we first remap 2D unwrapped coordinates back to reference Cartesian 3D coordinates. Between each time-point, the distance is multiplied by the isotropic scale given by the temporal rigid-body registration from STrEAMS to account for global embryo growth during this time interval.

#### Total curvilinear speed, v_total_

The sum of the magnitude of the 3D displacement vector, 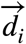 moved on the curved embryo surface between each timepoint *i*, over the total duration, *T*.

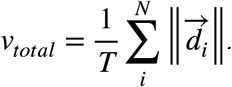

#### Mean curvilinear speed, v_mean_

The magnitude of the resultant sum of the 3D displacement vector, 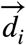, moved on the curved embryo surface between each timepoint *i*, over the total duration, *T*.

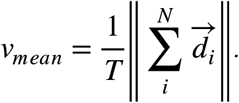

It corresponds to measuring the direct curvilinear distance travelled between the initial and final surface coordinates along a track.

#### Anterior curvilinear velocity, v_ant_

The resultant sum of the anteriorwards projection of the 3D displacement vector, 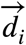 moved on the curved embryo surface between each timepoint *i*, over the total duration, *T*.

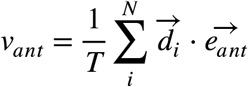

where 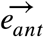 denotes the local anteriorwards unit surface vector and ± is the scalar dot product. 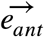 is computed by finding the equivalent 3D surface vectors when the vertical ‘upward’ anterior 2D vector in the unwrapped 2D polar projections is mapped back onto the reference embryo surface (Figure S4) (Stower et al. 2023). *v*_*ant*_ is positive if there is net movement anteriorwards, and negative if there is net movement posteriorwards.

### Tissue deformation analysis

To quantify the extent of how cell motion effects the dynamics of surrounding cells – the tissue deformation, in the VE and epiblast, we assessed the extent a control mesh grid was deformed by the resultant cell motion during the different DVE migration stages. Two metrics were measured to quantify deformation; the rate of change in i) surface area, and the rate of change in ii) curl (rotational) strain rate. The deformation analysis are conducted for VE and epiblast unwrapped projections independently. For two-channel videos, the epiblast expressing membrane-targeted tdTomato channel which simultaneously captures VE and epiblast motion comparable to Lifeact-GFP was used for motion extraction and deformation analysis.

#### Construction of the fine-grained 640 sector control grid from the coarse 32 sectors used for statistical analysis

A 32 sector grid was subdivided equally by 5 angular and 4 distance intervals such that each sector is tessellated without overlap by 5×4=20 smaller quadrant ROIs. These subdivision parameters were chosen so that the sub-sectors were a good quadrilateral approximation (same surface area) of the continuous corresponding curvilinear surface patch in Cartesian 3D. Each ROI region is modelled as a quadrilateral in 2D by linearly joining its 4 corner points; (*i*_1_, *j*_1_), (*i*_2_, *j*_2_), (*i*_3_, *j*_3_), (*i*_4_, *j*_4_) and correspondingly in 3D where (*i*_1_, *j*_1_) ⟷ (*x*(*i*_1_, *j*_1_), *y*(*i*_1_, *j*_1_), *z*(*i*_1_, *j*_1_)) etc. when reversing the unwrapping transform.

#### Deformation of the fine-grained 640 sector control grid by cellular motion optical flow

The corner points of each of the 640 ROI grids was propagated frame to frame using forwards Euler;

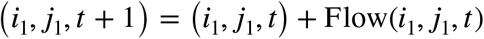

where *t* is the frame index, until the end of the respective migration stage and Flow(*i, j, t* is the MOSES extracted DeepFlow optical flow from the VE and epiblast unwrapped 2D projection videos.

#### Measurement of rate of change in surface area

The mean fractional rate of change in surface area, 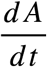 of a local tissue patch was measured as previously reported (Stower et al. 2023) where 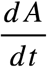 at time *t* is defined as 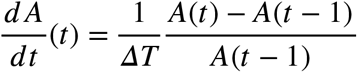 where A = Area(t)/Area(t-1), and *ΔT =* 5 min for Lifeact-GFP and *ΔT =* 10 min for Hex-GFP:mem-tdTomato and Ttr-Cre:ROSA26^mTmG^ embryos, the time elapsed between individual frames and 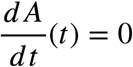 for time *t* = 0. The control grid coordinates were mapped back to true Cartesian 3D coordinates and the 3D quadrilateral surface area at time *t*, was computed as

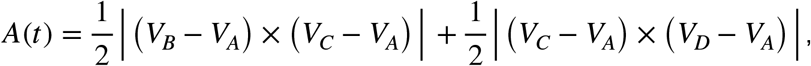

the sum of the area of the two constituent planar triangles △ *ABC* and △ *ACD* where *V*_*A*_, *V*_*B*_, *V*_*C*_, *V*_*D*_ are the corresponding vertex (*x, y, z*) coordinates of the quadrilateral *ABCD* . For visualisation, we smoothed the 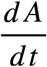 for each of the 640 control grid tissue surface sectors temporally, running asymmetric least squares (Eilers and Boelens 2005) with parameters *p* = 0.5, *λ* = 100 for 10 iterations after edge-mode padding of signals by the maximum of *T*/100 or 3 frames.

#### Measurement of the rate of change in isotropic, anisotropic and curl strain

This analysis assesses the extent of deformation of the control grid through measuring the rate of change of the mesh links or edges. We show below this is equivalent to monitoring the velocity gradient, which can be further decomposed into independent contributions due to isotropic (expansion/dilation), anisotropic (shear) and curl (rotational) strain rates. The analysis assumes cellular velocities in the tissue are small between time points and vary smoothly across the embryo, which are reasonable assumptions for the tissue motion observed in the VE and epiblast.

For each 2D sector in the control grid at time *i*, we remap its 2D (*i, j, t*) corner points back to Cartesian 3D to get the corresponding 3D surface coordinates, (*x*(*i, j, t*), *y*(*i, j, t*), *z*(*i, j, t*)).

Maintaining the same connectivity, this generates the corresponding 3D sector. For a 3D sector, we define the edge or link *l*, as the displacement between a pair of vertices, *r*_1_ = (*x*_1_, *y*_1_, *z*_1_) and *r*_2_ = (*x*_2_, *y*_2_, *z*_2_).

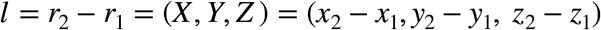

The rate of change of the edge is the relative velocity between the two vertices, 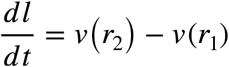.

If the local tissue motion is smoothly varying and small between timepoints, we can use an affine approximation and linearly interpolate the velocity at two vertices, *v*(*r*_2_) ≃ *v*(*r*_1_) + ∇*v*^*T*^ ⋅ (*r*_2_ − *r*_1_.

The rate of change in link length can thus be expressed in terms of the velocity gradient, ∇*v*.

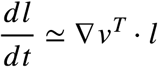

This expression can be rearranged to give an expression for ∇*v* in terms of *l* by by taking the right outer product with *l*,

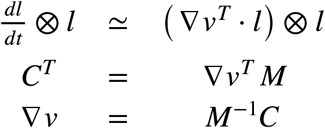

where 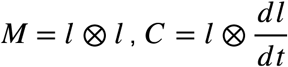 and ⊗ denotes the tensor outer product. Letting *W* = *M*^−1^*C*, we proceed to decompose ∇*v* = *W* = *W*_sym_ + *W*_anti−sym_ uniquely into a symmetric, 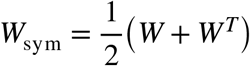 and anti-symmetric component, 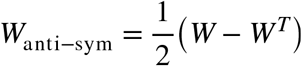.

The symmetric component, *W*_sym_ is the linearised continuous total strain rate defined as 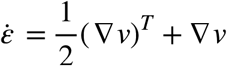 for infinitesimal deformations of a continuum body. As it is a symmetric matrix, it can further be decomposed *W*_*sym*_ = *W*_*iso*_ + *W*_*aniso*_ into an isotropic (scaled-identity matrix), 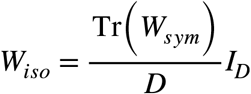 and an anisotropic (off-diagonal only matrix), *W*_*aniso*_ = *W*_*sym*_ − *W*_*iso*_ component where Tr(⋅) denotes the matrix trace and *I*_*D*_ is the identity matrix in dimension *D*. Eigenvector analysis can further be applied to *W*_*aniso*_ to quantify the strain ratio in principal strain directions (Rozbicki et al. 2015).

The anti-symmetric component, *W*_anti−sym_ measures the local rotation rate of tissue motion or curl, *W*_*curl*_. In summary, we have the following complete decomposition of the velocity gradient.

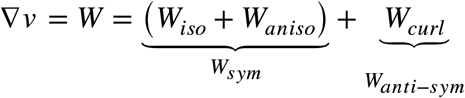

In practice, the computations described were not implemented in Cartesian 3D. The sectors are local surface elements therefore for improved numerical stability we defined a local 2D surface coordinate system based on anterior-posterior and left-right directions. The edge vectors were projected with respect to this 2D basis which reduces the above computations to 2D. For visualisation, we smoothed the measured strain for each of the 640 control grid tissue surface patches temporally, running asymmetric least squares smoothing with parameters for 10 iterations after edge-mode padding of signals by the maximum of T/10 or 3 frames.

#### Averaging measurements from 640 ROI fine-grained sectors to the 32 analysis sectors

For a migration stage of total duration *T*, each deformation metric is measured in each of the 640 ROIs as a time-series of length *T* − 1. To compute the average metric for the 32 sector analysis, the mean temporal value of the metric is computed first for each time-series, then the mean is computed over the mean temporal values of the 20 component sub-sectors in each of the 32 sectors.

### DVE cell tracking – directionality analysis

This analysis is performed on 5 Lifeact-GFP embryos, with single cell tracking (Stower et al. 2023). For a given time interval, we compute the weighted angle histogram considering all cells located in the distal tip sector region-of-interests (ROIs) i.e. sectors 1-8, in each constituent timepoint within the interval, that is moving (speed > 0). The histogram partitions the angular space of ±180^0^ relative to the A-P axis into 16 bins. Cell velocity, *v* = (Δ*x*, Δ*y*) is measured in 2D geodesic projections as the difference between its current (*x* (*t*), *y*(*t*)) centroid position and that in the next timepoint ;(*x* (*t* + 1), *y*(*t* + 1)), (Δ*x*, Δ*y*) = (*x* (*t* + 1), *y* (*t* + 1)) − (*x* (*t*), *y* (*t*)). The cell’s angle at time *t* is 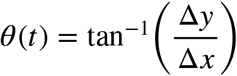 and its speed, 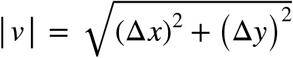. The weighted angle histogram allocates each cell to an angular bin according to *θ* (*t*), with an associated weighted ‘count’ of |*v*| . The Rayleigh statistic, 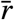 is computed for each time interval based on all *N* individual cell instances, 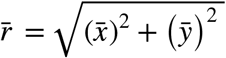 where 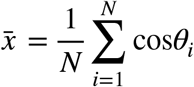 and 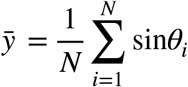. We compute the histogram for 5 continuous time intervals, each 10 min (2 timepoints) for each of pre-migration/initiation, and migration phases, for each embryo. The average embryo histogram is the mean of the sum-normalised individual embryo histograms.

### Extended depth apical-to-basal VE and epiblast polar cartographic projections

We extend the VE and epiblast surface used in polar Cartographic projections of the registered embryos, by three sets of surface propagation:

1. Outward propagation of unwrapped VE surface to cover the entirety of any VE cells.
2. Linear interpolation of unwrapped VE surface to its aligned unwrapped epiblast surface
3. Inward propagation of epiblast surface to cavity.

The first and third surface propagation are implemented by forward Euler updates with a step-size of 1 voxel along the gradient of the Euclidean distance transform (EDT) of the corresponding binary volumes of the unwrapped VE and epiblast surfaces. To ensure correct propagation at the ExE end of the embryo, binary volumes were concatenated with a ‘mirrored’ version prior to computing EDT. Interpolating the image intensity at the propagated surface coordinates generates the extended depth polar Cartographic projection.

### Extended depth midline sagittal projection

The midline sagittal projection image and unwrapping coordinates resamples those of the extended depth apical-to-basal VE and epiblast polar Cartographic projection, along the A-P axis.

### Analysis of ratchet-like motion in Hex-GFP labelled DVE cells

For each Hex-GFP:mem-tdTomato embryo, we computed its extended depth mid-line sagittal projection videos and applied MOSES to the mem-tdTomato channel to compute the frame-to-frame optical flow motion jointly for VE and epiblast cells. We thresholded the Hex-GFP intensity, and retain the largest contiguous spatial component as the DVE cell population segmentation in each frame. Then we average the 3D area-distortion corrected optical flow with the DVE cell binary segmentation to compute the mean DVE velocity in each frame – the DVE velocity time-series. In the midline sagittal unwrap, the x-axis is the A-P axis, therefore the x-component of DVE velocity time-series is the anteriorwards velocity time-series. To assess ratchet-like motion, we computed (i) the fraction of time-points across all embryos with positive anteriorwards velocity, (ii) verifying the autocorrelation of the anteriorwards velocity is oscillatory by detecting peaks and extracting the mean peak-to-peak temporal difference.

### Midline kymograph visualisation of 2D polar projections and associated statistics

Single cell associated statistics were mapped to a pixel-based 2D polar geodesic images as described above for “visualisation of cell statistics in 2D projections”. The boundaries of the sectors were propagated as described above for tissue deformation over the video duration and rasterised to a binary 2D projection time-lapse. Other data such as image intensity, MOSES extracted optical flow and local embryo shape change displacements are already computed in the format of pixel-based 2D polar geodesic images. A kymograph of the data of interest is visualised by concatenating the values at all pixels along the aligned A-P axis over time.

### Visualisation of cell statistics on 3D meshes

To visualise the desired scalar statistics of a segmented cell instance in 3D we used MeshLab (Cignoni et al. 2008). Meshlab requires as input a triangle mesh (trimesh). We use Python *Trimesh* library to write a .obj trimesh which requires a list of 3D (*x, y, z*) vertex coordinates, a list of faces, and a 3-tuple specifying how vertex indices are connected together in triangles and a list of vertex colours, and a 3-tuple specifying the RGB colour at each vertex. Cell statistics were first visualised in the 2D projections as an image as described above with cell tracks drawn as 3-pixel wide lines using Python *Scikit-Image*. A 2D image is equivalent to a quadrilateral mesh specified by a list of 2D (*i, j*) vertex coordinates, a list of quadrilateral faces, 4-tuples specifying how individual pixels are connected to neighbours in squares and the 2D image pixel colour as the RGB vertex colour. To obtain the desired 3D trimesh, we triangulate the quadrilateral mesh to a trimesh using the Python *Trimesh* library (*trimesh*.*geometry*.*triangulate_quads* function); remove all (*i, j*) vertex coordinates and faces not part of the VE in the polar geodesic projection; and remap the 2D (*i, j*) coordinate to 3D (*x, y, z*) vertex coordinates using the 2D-to-3D unwrapping coordinates (see above).

### Vertex model for DVE migration

We model the tissue using a 2D vertex model (Alt et al. 2017; Fletcher et al. 2014) where each cell is represented as a polygon with vertices as the degrees of freedom. The equation of motion of the vertex is given by the following overdamped dynamics in the vertex model

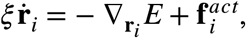

where *ξ* is the substrate friction coefficient and 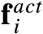 is an active force on vertex *i*. 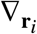 is the gradient of the energy function *E* evaluated with respect to the position vector of vertex *i*. The energy function 45 is

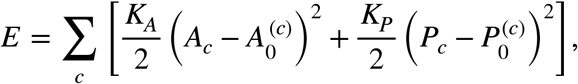

where the summation is over all cells. *A*_*c*_ and *P*_*c*_ are the area and perimeter of cell 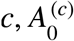 and 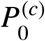 are the target area and perimeter for cell *c*, and *K*_*A*_ and *K*_*P*_ are the area and perimeter elasticity moduli. The quantity 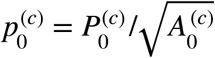 defines a target shape index. It is well known that a passive vertex model (with all cells having same target area and target perimeter) undergoes a rigidity transition near 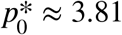 (Bi et al. 2015). Hence for 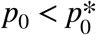 the tissue behaves like a solid whereas for 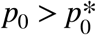 the tissue behaves like a fluid. We fix *K*_*A*_ = 1 *K*_*P*_ = 0.02 *ξ* = 1.

We model DVE migration by considering three different cell types in the vertex model:

1. *DVE* cells. In the geometry of the setup, DVE cells form a small cluster of *N*_*DVE*_ = 47 located near the centre of the tissue. DVE cells are “active” with a self-propulsion force, *F*, in the positive vertical direction + *y* i.e., along the anterior-posterior axis. In experiments, these cells are observed to have smaller apical area than surrounding cells. Hence, we gives these cell a smaller target area 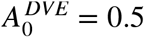 compared to the surrounding (emVE and exVE) cells. Further, in experiments these cells are observed to have higher levels of apical tension and F-actin keeping them together as a tight cluster, so we fix their target perimeter to 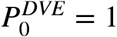, so that their target shape index 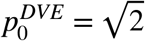 is well below the rigidity transition and these cells act as an effective solid-like cluster.
2. *emVE* cells. These are passive cells with their target perimeter 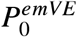 and target area 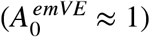 larger than the values of DVE cells. The target shape index 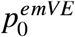 for these cells is still below the rigidity transition. While these cells are therefore solid-like, they can still rearrange if stresses in the tissue are sufficient to overcome the energy barriers for T1 transitions.
3. *exVE*. These cells form the outer layer of the 2D tissue in our model and they mark the end of DVE migration. These cells in experiments act as a “wall” for the migrating DVE cluster. Hence, in the simulation, the vertices of these cells are not allowed to move at all and they form a boundary marking the end of DVE migration.

Our 2D vertex model consists of an 18 × 18 grid of cells with a total of *N* = *N*_*DVE*_ + *N*_*emVE*_ + *N*_*exVE*_ = 324 cells. We fix 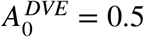 for DVE cells while 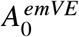 is selected such that the mean target area: 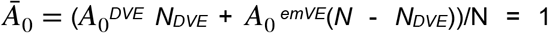. For DVE cells we have the target shape index as 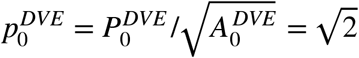 whereas for emVE cells we have 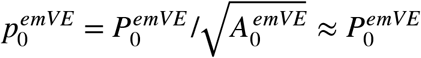 since 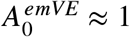. Our chosen parameters sets the units of length as 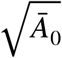 *and time as ξ* /(*K*_*A*_ *Ā*_0_).

We simulate the system by first creating a disordered tissue from a regular hexagonal lattice. This is done by adding tension fluctuations in the cell-cell junctions (Curran et al. 2017; Krajnc 2020). Once the tissue has been disordered, DVE, emVE, and exVE cells are selected and assigned their target areas and perimeters (exVE use the same values as emVE cells), and the tissue is allowed to relax for a duration of 1000 time units. This tissue forms the initial configuration of the system and exVE cells are fixed at this point. From here, an active force is applied to the DVE cluster that acts in the + *y* direction and is ramped up linearly from 0 to *F* from *t = 0* to *t = 2000*. Beyond this time, the active force is held at a constant value of *F* for the remainder of the simulation (*t* = 2000 to *t* = 10000).

Equations of motion are solved using an explicit Euler scheme with time step 0.01. Edges that fall below a threshold length 0.01 undergo a T1 transition, with final length set to 0.011 (if an edge contains a ‘fixed’ vertex belonging to an exVE cell, it cannot undergo a T1). To add a self propulsion force on each DVE cell, we follow the prescription in (Sussman 2017). Each cell is assigned a self propulsion force *α*_*i*_. The additional active force on each vertex then reads

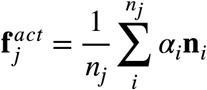

where the sum is over all *n*_*j*_ cells that share vertex *j* and **n**_*i*_ = (0,1) is a unit vector along the direction of the polarity of cell . We set *α*_*i*_ = 0 for the passive cells, whereas for the DVE cell *α*_*i*_ goes linearly from 0 to *F* between *t = 0* and *t = 2000* and then remains constant.

The *y* component of the mean velocity of the DVE cluster is determined by first calculating the *y* component of the mean position of the DVE cluster by averaging over the coordinates of the centre (mean of vertex positions) of each cell that belongs to the DVE cluster. Following this, a finite difference in time is used to calculate the time-average velocity, i.e., average change in *y* position of the DVE in a given time.

In order to distinguish the regime of intermittent migration from that of more smooth migration that does not have velocity bursts (Figure 5B,B’), we plot the maximum value of the *y* component of DVE mean velocity and its coefficient of variation. Both of these quantities were calculated by analysing the velocity time series of the *y* component of the DVE mean velocity from *t* = 3500 to *t* = 10000. The initialisation of the analysis at allows sufficient time for (i) DVE clusters that do not migrate completely to come to a halt, and (ii) DVE clusters that migrate relatively smoothly to reach the emVE-exVE boundary and come to a stop. This allows us to identify a strip in the parameter space that corresponds to the intermittent migration of the DVE cluster. The coefficient of variation (CV) is the standard deviation of the *y* component of DVE mean velocity divided by its mean value, calculated in the interval *t* = 3500 to *t* = 10000. Note that, CV is only calculated if the mean value is sufficiently large (i.e. greater than 10^−5^). We expect that for intermittent migration, fluctuations in the velocity time series are larger than the mean value, hence regions corresponding to CV greater than 1. Furthermore, the intermittent regime in the parameter space also approximately corresponds to large maximum DVE velocity between *t =* 3500 and *t =* 10000. Hence, large values of maximum DVE velocity and CV in Figure 5B’-B’’ correlate with regions of intermittent DVE migration in the parameter space.

## Supplemental Data Figures and Tables

**Figure S1.**
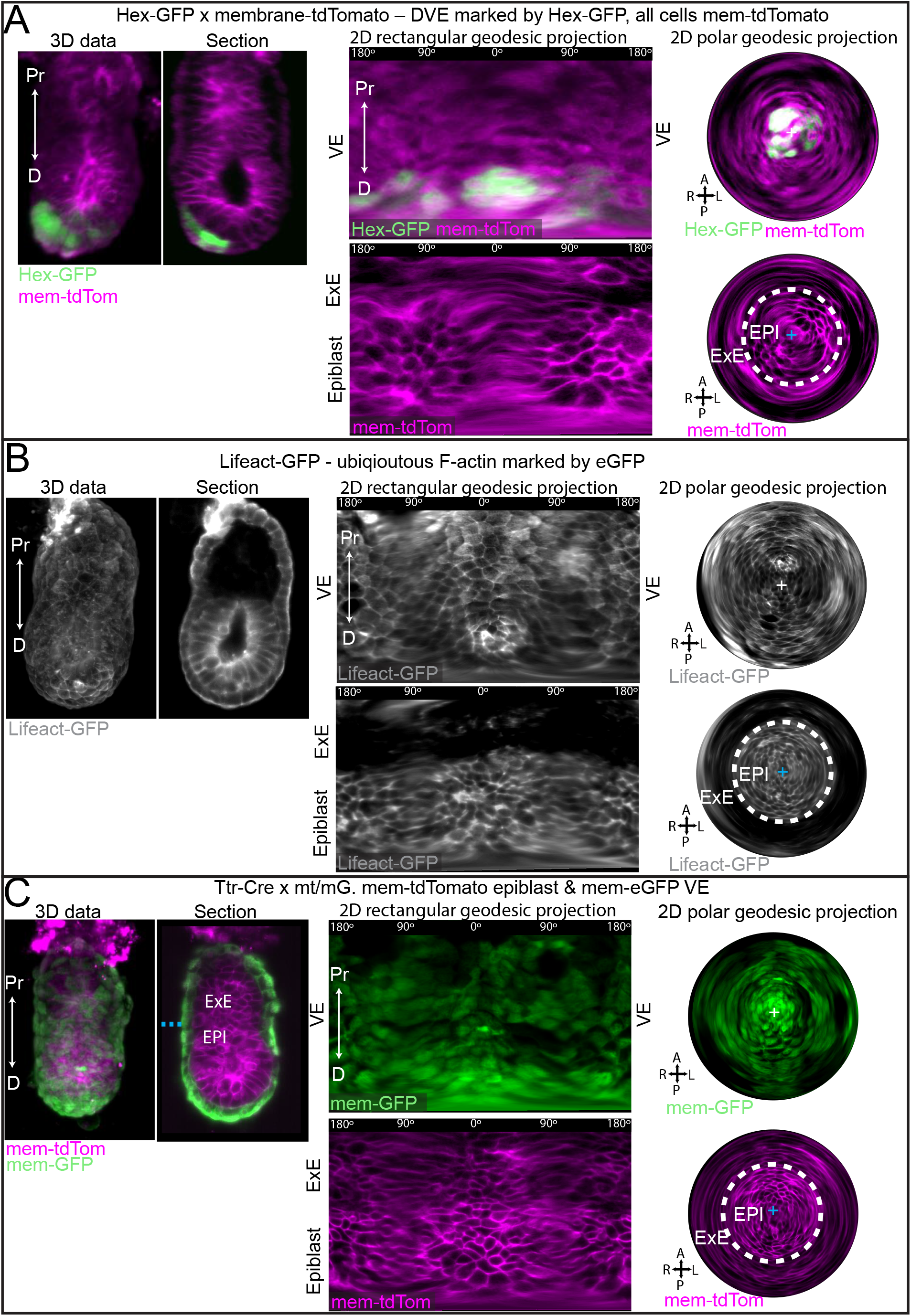
Examples of light-sheet time-lapse data and geodesic projections of VE and epiblast tissues in three fluorescent reporter mouse lines. **(A)** Example Hex-GFP; membrane td-tomato, **(B)** Lifeact-GFP, and **(C)** mTmG crossed with Ttr-Cre line. Each panel shows a max intensity projection, mid-sagittal optical section, and rectangular and polar projections of the apical surface of the VE (upper panels) and basal surface of the epiblast (lower panels). We note that the Lifeact-GFP mouse line consistently showed an almost complete absence of fluorescence signal in the ExE tissue. This does not reflect the level of F-actin in the ExE as shown by immunofluorescent staining. Note projections have no scale bar as they are non-linear.

**Figure S2.**
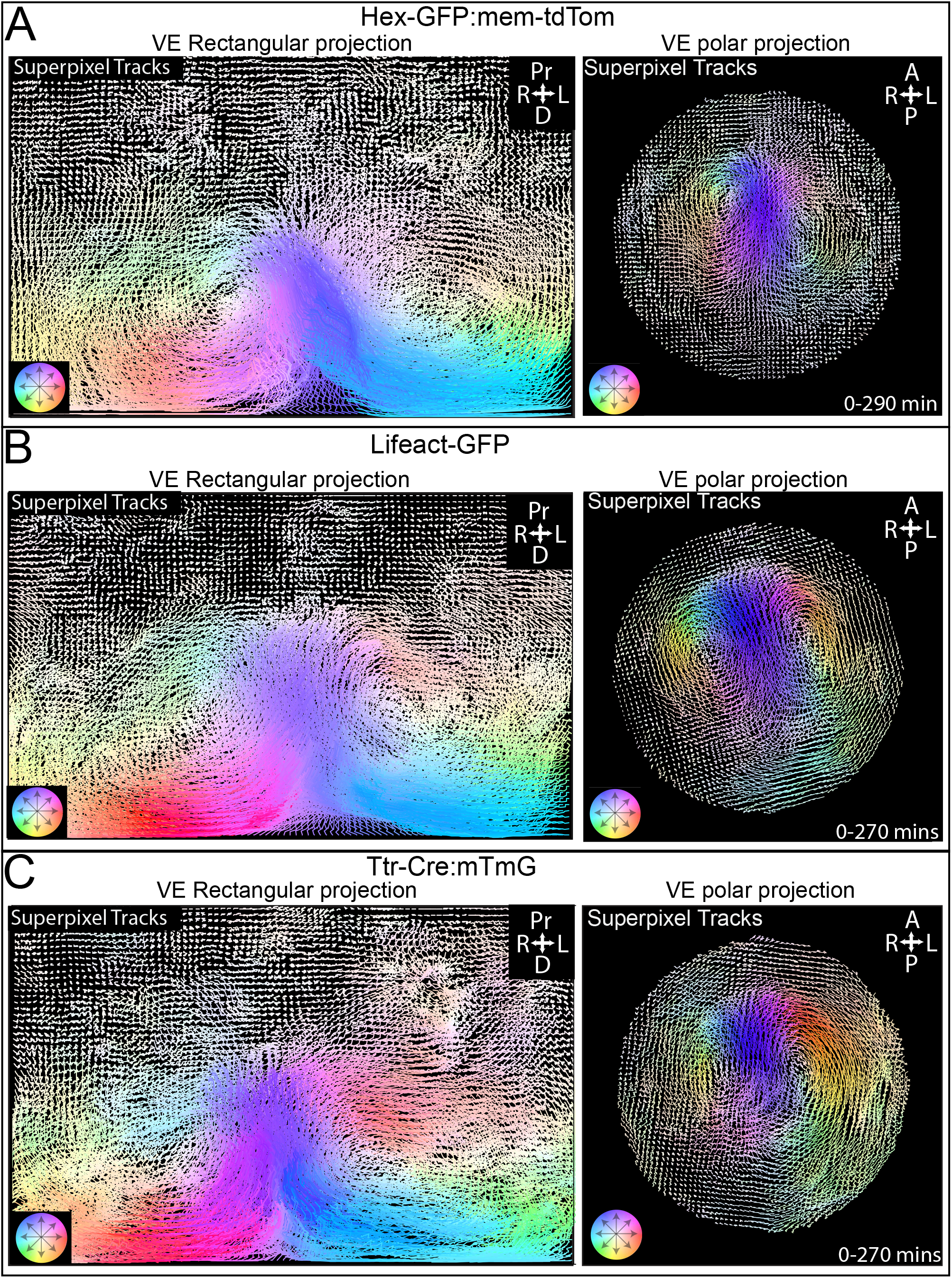
Superpixel motion tracking of the VE tissue on geodesic projections from three fluorescent mouse lines. **(A)** Example Hex-GFP; membrane td-tomato, **(B)** Lifeact-GFP, and **(C)** mTmG crossed with Ttr-Cre line. Each panel shows superpixel tracks on a rectangular geodesic projection and polar geodesic projections of the apical surface of the VE. Tracks are coloured by the direction of motion. Note re-projections have no scale bar as they are non-linear.

**Figure S3.**
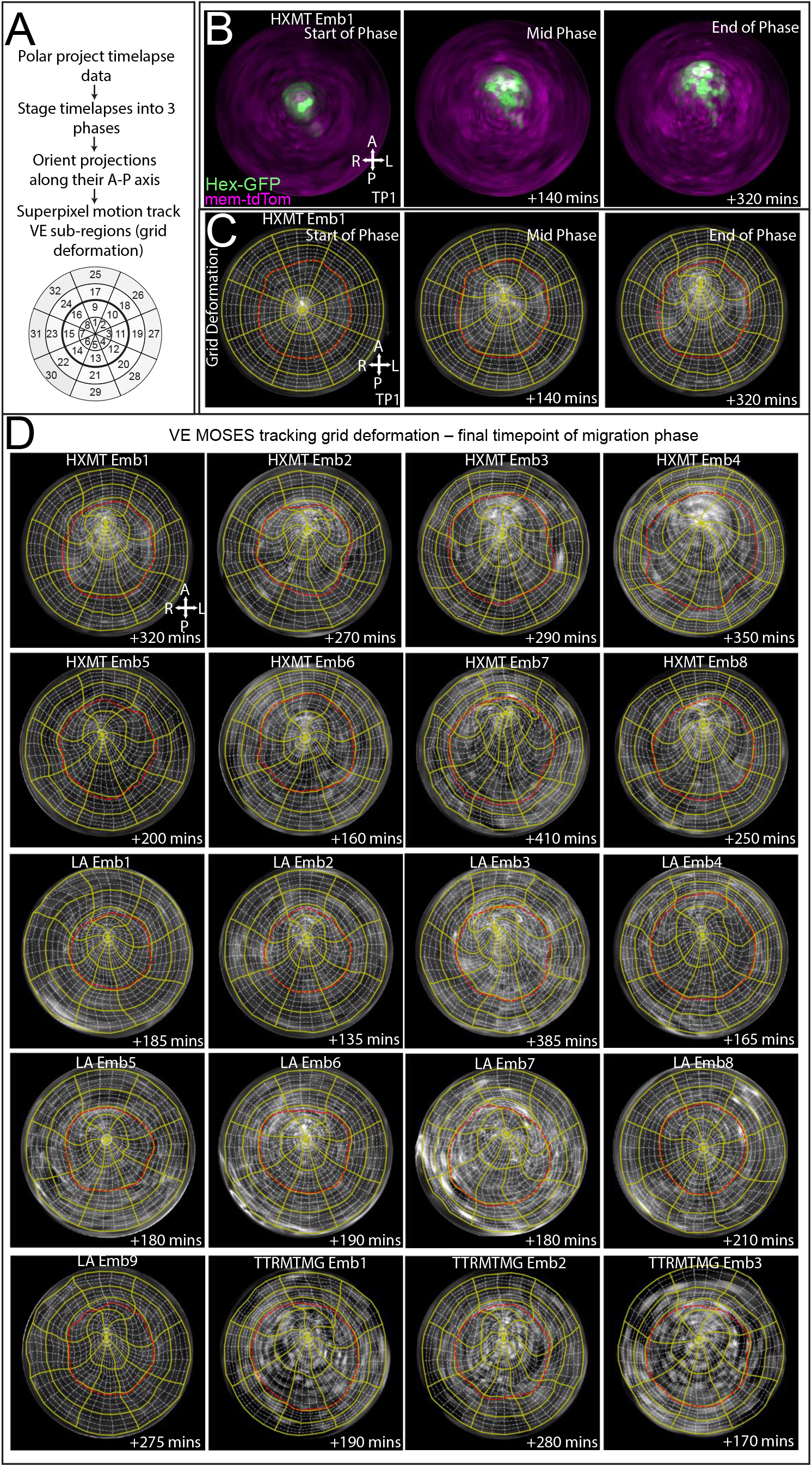
Sub-regional tracking of the VE tissue using MOSES superpixel motion tracking. **(A)** Overview of method for tracking the motion of VE sub-regions using superpixel motion tracking. **(B)** Example Hex-GFP;mem-tdTomato embryo showing the start, middle and end of migration in a geodesic polar projection. **(C)** Superpixel motion tracking of embryo in B. As the tissue moves the grid is deformed, enabling analysis of the tissue behaviour within each tissue sub-region and comparison across embryos. **(D)** Final timepoint of superpixel motion tracking from all (n = 21) embryos in this dataset. All data is converted back to 3D coordinates for analysis. For each stage (pre-migration/initiation, migration, post-boundary) the grid is re-set and re-tracked by motion behaviour. Note re-projections have no scale bar as they are non-linear. HXMT = Hex-GFP;membrane-tdTomato, LA = Lifeact-GFP, TTRmTMG = mTmG;Ttr-Cre embryo.

**Figure S4.**
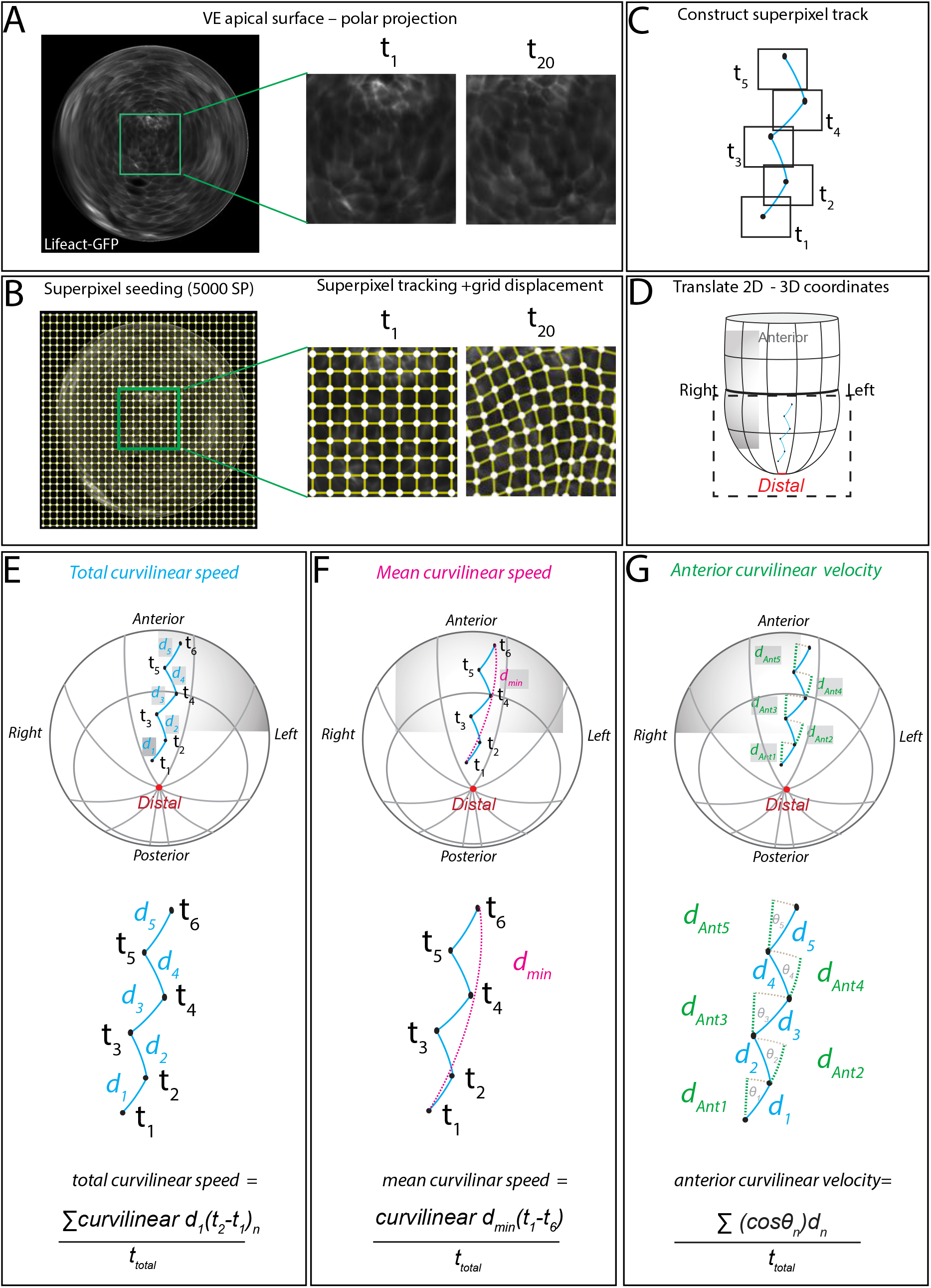
Overview of analysis of motion superpixel tracks from time-lapse data. **(A)** Example Lifeact-GFP embryo reprojected as a geodesic polar projection from 4D light-sheet imaging. **(B)** A uniform grid is seeded at the start of each migration phase and the motion in each sub-region tracked throughout the time-lapse. **(C)** Superpixel tracks are constructed for each region. **(D)** The apical surface coordinates of each track are converted back to 3D coordinate space. **(E-G)** Three parameters of motion behaviour calculated for each superpixel track: total speed, mean speed and anteriorwards velocity.

**Figure S5.**
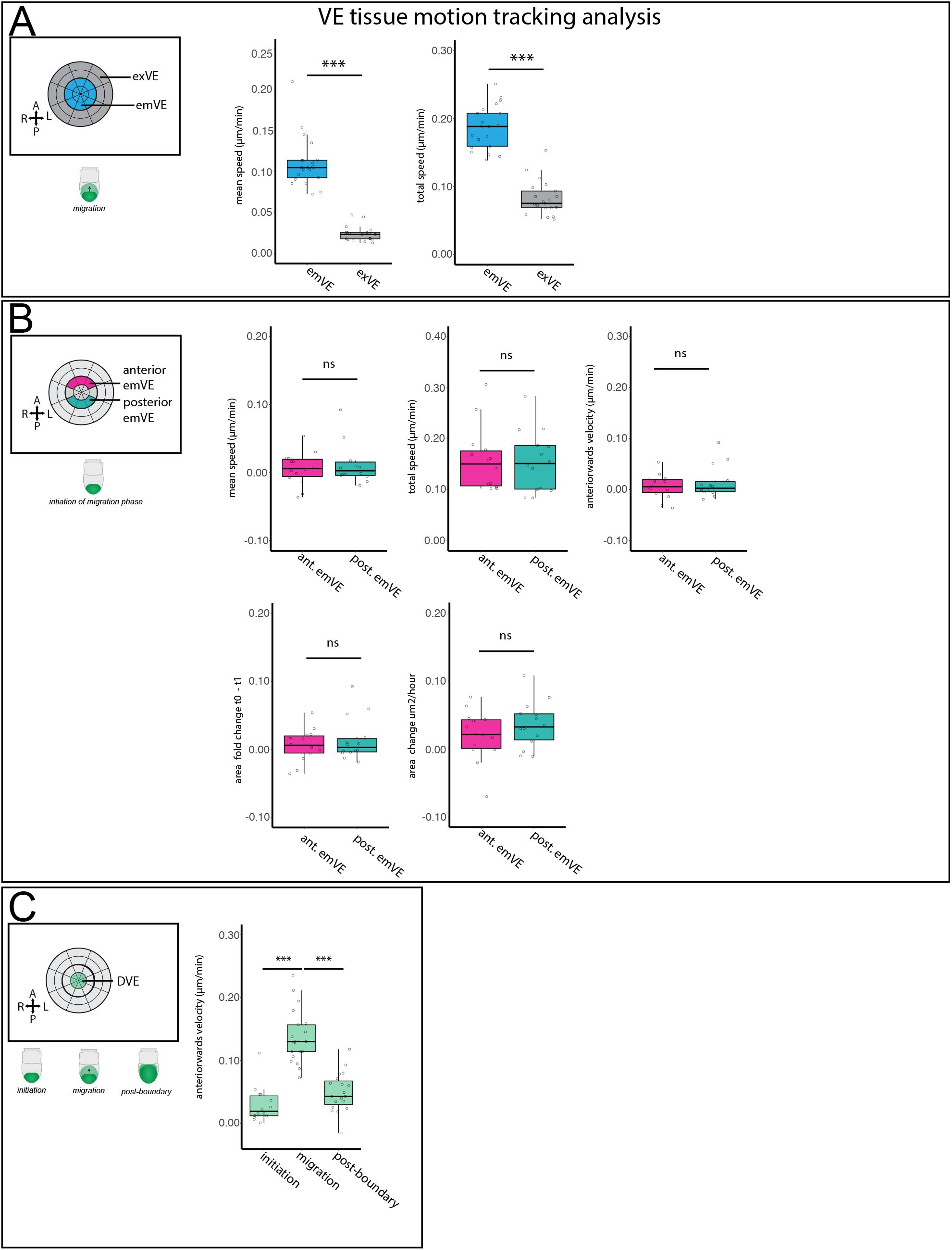
Analysis of MOSES superpixel tracking of VE tissue. **(A)** Superpixel motion tracking of the emVE and exVE tissues confirms that emVE is significantly more active (higher mean speed and total speed) than the exVE (for both comparisons; Student’s *t*-test, *p*=<0.001). **(B)** Comparison of the anterior emVE and posterior emVE that are immediately adjacent to the DVE during the initiation of migration phase. There is no significant difference in motion (mean speed, total speed or anteriorwards velocity) or tissue area change (area fold change or area rate change) between these two tissue regions (all comparisons; Student’s *t*-test, *p*=>0.05). **(C)** Superpixel tracking of the distal VE during each phase shows that the migration phase has the highest anteriorwards velocity (one-way ANOVA, *p*=<0.01, followed by Tukey’s HSD Test on initiation of migration vs. migration (p =<0.001), migration vs post-boundary (*p*=<0.001), initiation of migration vs post-boundary (*p*=>0.05)). In the post-boundary phase the distal VE region contains the nascent Hex-GFP cells that continue to migrate anteriorly, but do so at a lower speed than the initial DVE.

**Figure S6.**
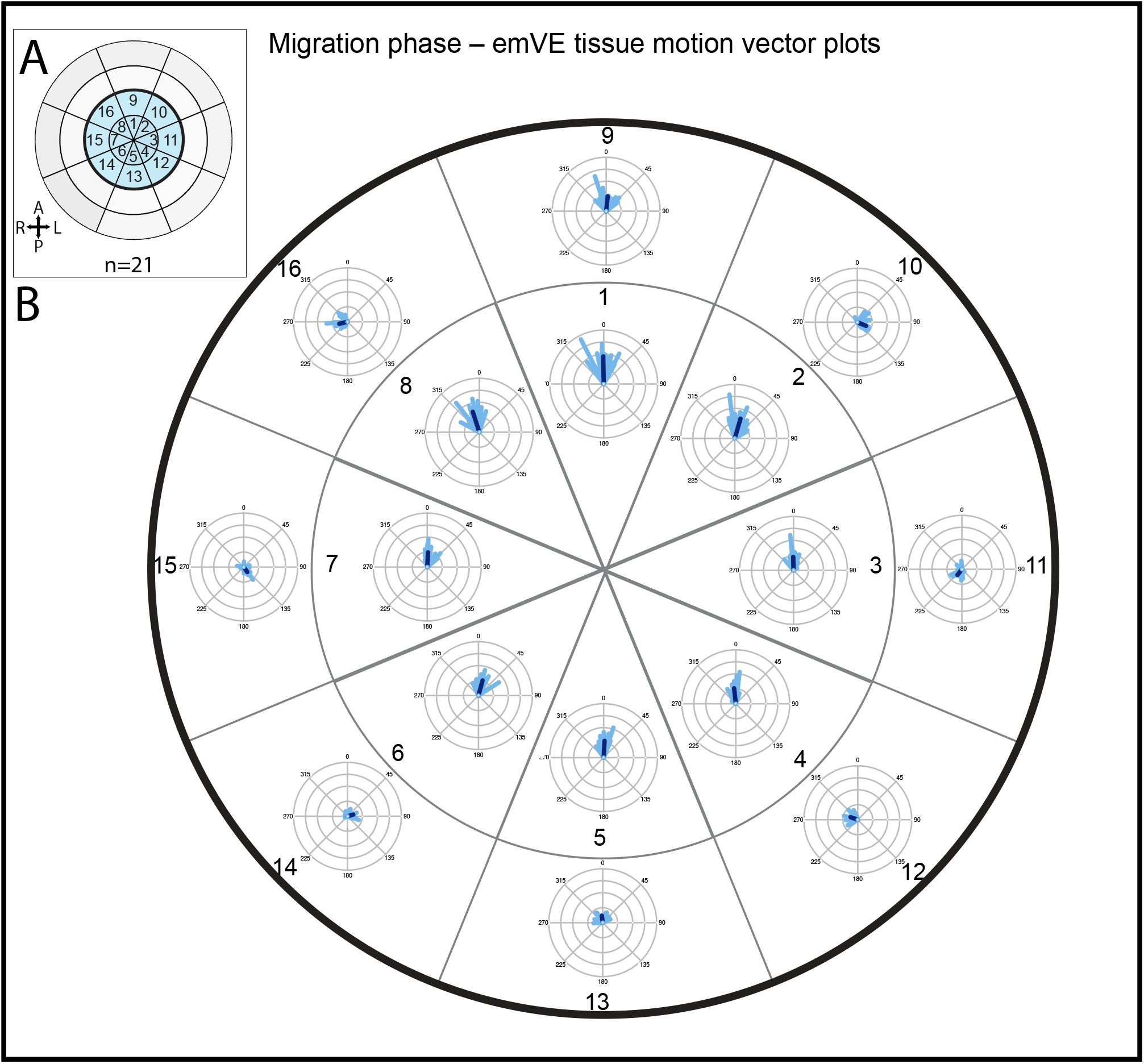
Motion vector distribution in emVE during DVE migration. **(A)** Diagram of polar view of VE. emVE is the central, blue coloured region. **(B)** For each sub-region of the emVE the median motion vector of superpixel tracks is calculated and plotted as a windrose plot for a total of 21 embryos. While the DVE region (sectors 1-8) show highly aligned anteriorwards vectors, there is a bi-rotational flow around the emVE (sectors 9-16). The length of lines is plotted by average superpixel velocity for each region – note the longer lines in the DVE region (sectors 1-8) compared to those of the surrounding emVE (sectors 9-16). Analysis of the distribution of vector angles (see Table S2 for Rayleigh distribution statistic) shows a significant Rayleigh distribution statistic across each emVE sub-region, showing this pattern of motion is conserved across embryos.

**Figure S7.**
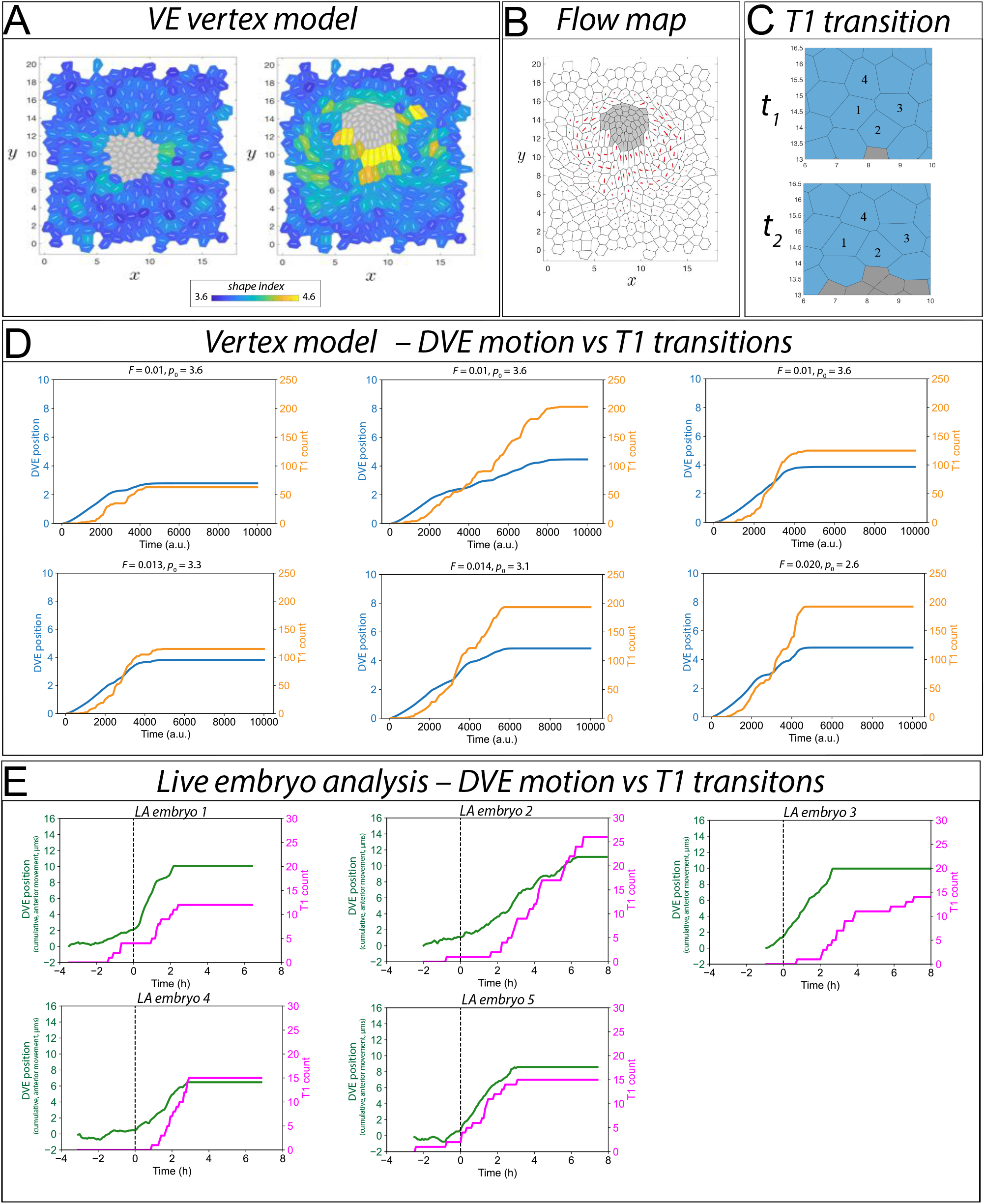
Analysis of ratchet-like DVE movement and T1 transitions *in silico* and *in vivo*. **(A)** Example simulation at *t* = 0 and *t* = 6000 with cells coloured by shape index. **(B)** Flow maps of cells in simulation in A show a similar rotational flow to the mouse visceral endoderm. **(C)** Example T1 transition in simulation showing the intercalation of cell 2 between 1 and 3. **(D)** 6 example vertex simulations varying DVE self propulsive force, *F*, from 0.01 - 0.02, and emVE target shape index, *p0*, from more rigid to less rigid; 2.6 - 3.6. Orange line; T1 count. Blue line; DVE motion. **(E)** DVE motion and T1 transitions in 5 Lifeact-GFP E5.5 embryos. Magenta line; T1 count. Green line; DVE motion. Dashed vertical line shows the onset of directional DVE migration in embryos.

**Figure S8.**
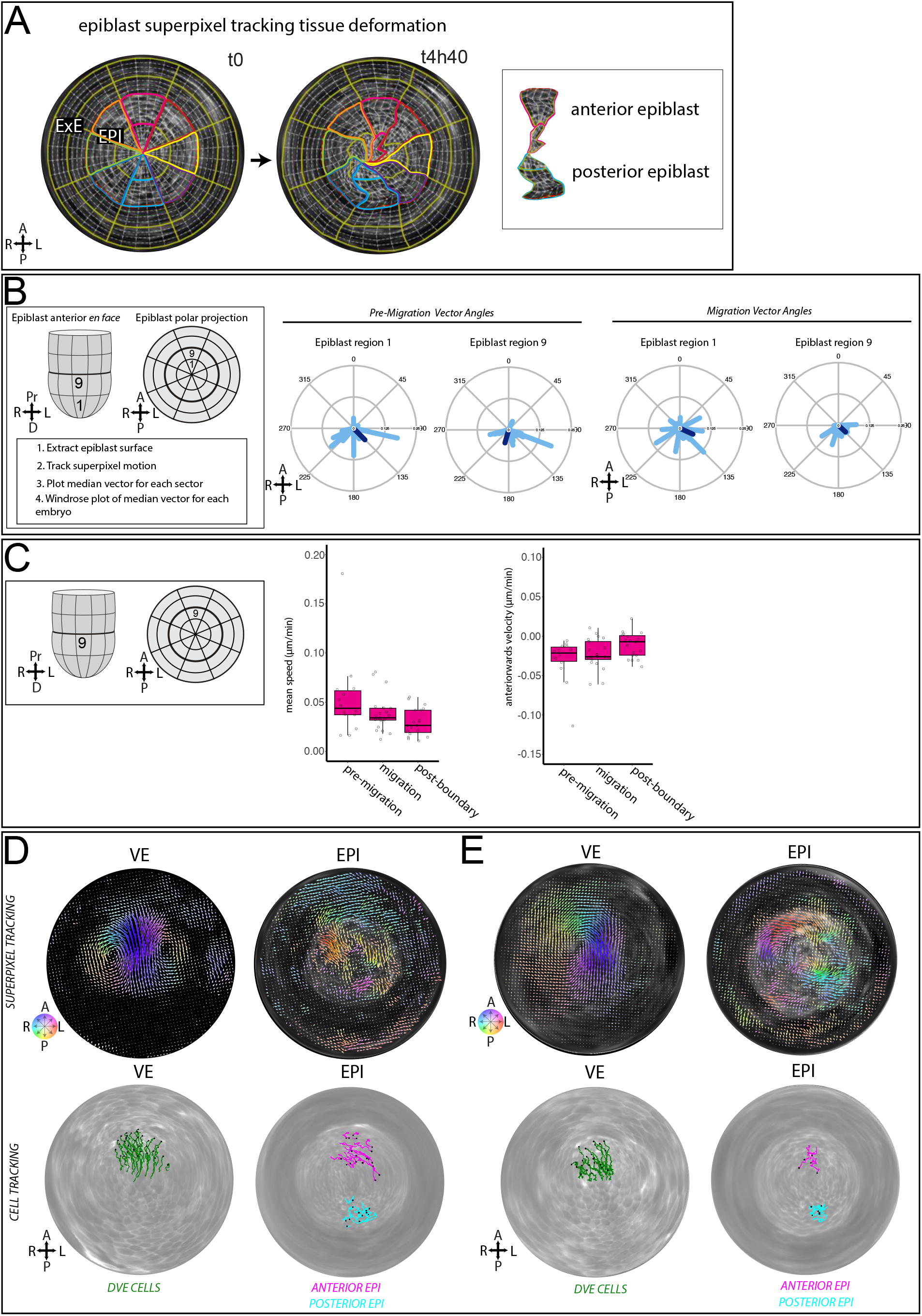
Motion analysis of epiblast tissue dynamics and epiblast cell tracking. **(A)** Example motion superpixel grid tracking of epiblast during DVE migration with anterior epiblast region (red) and posterior epiblast (blue) regions highlighted at the start and end of the phase. Inset showing anterior and posterior epiblast regions at the end of DVE migration. Anterior epiblast becomes elongated and shifts posteriorly. **(B)** Motion vector plots of anterior epiblast regions during DVE pre-migration/initiation and migration phases. **(C)** Analysis of mean speed and anteriorwards velocity components of superpixel tracking of anterior epiblast during migration phases. **(D**,**E)** VE and epiblast tracking of two example embryos. Top row shows superpixel tracking of VE and epiblast. Bottom row shows cell tracking of VE (green tracks), anterior epiblast (magenta tracks) and posterior epiblast (cyan tracks).

**Figure S9.**
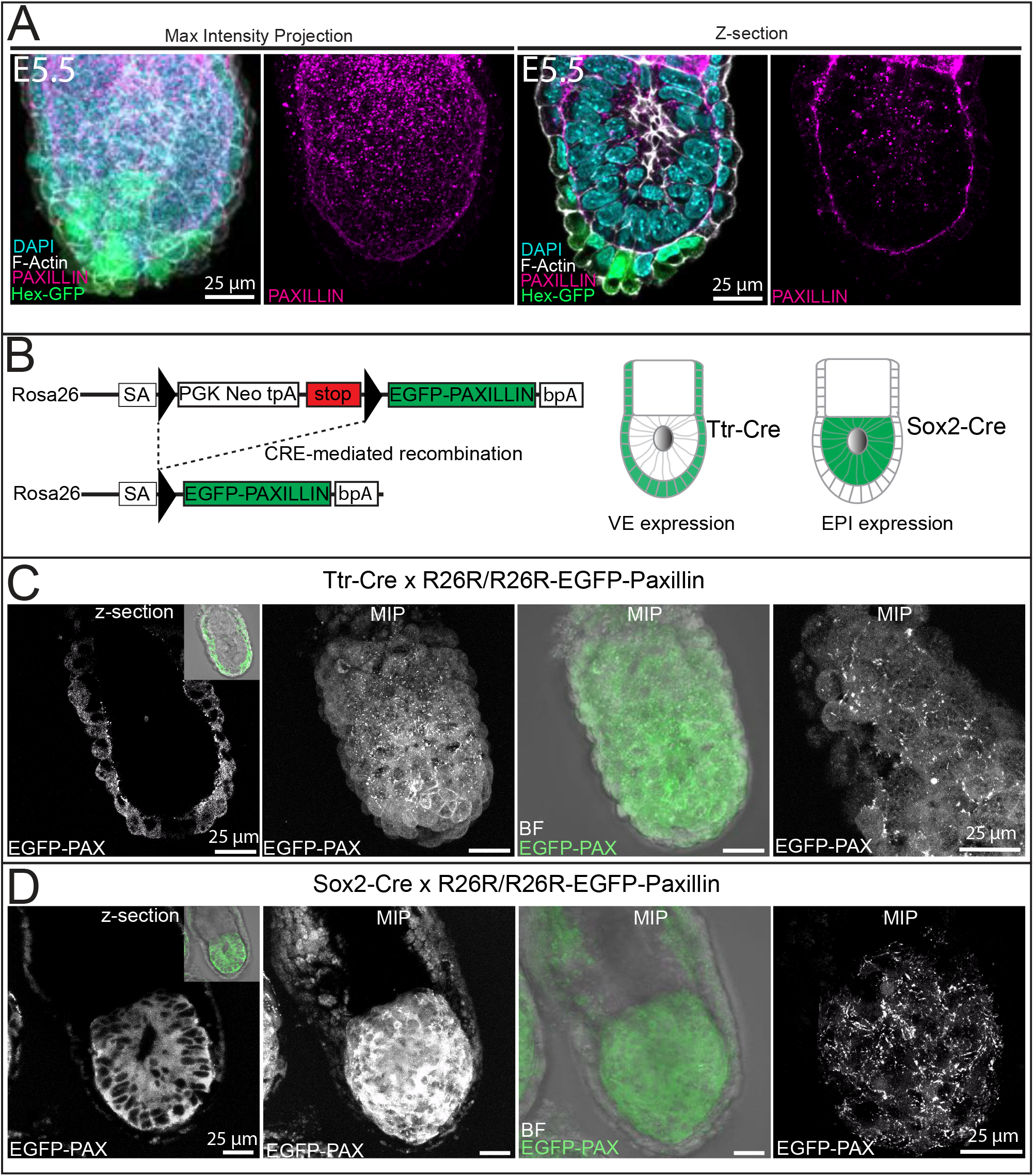
Paxillin localisation in the E5.5 embryo. **(A)** Whole-mount immunofluorescence of PAXILLIN in an example E5.5 embryo, showing a maximum intensity projection (MIP) and single z-section of the embryonic-half of the embryo. PAXILLIN is enriched along the basal surface of the VE and epiblast. **(B)** Diagram of Cre-inducible EGFP-PAXILLIN reporter line. Ttr-Cre is expressed specifically in the VE and Sox2-Cre in the epiblast. **(C)** Example live imaging of E5.5 Ttr-Cre EGFP-PAXILLIN embryo and **(D)** Sox2-Cre EGFP-PAXILLIN embryo, showing the basal localisation of the protein throughout the VE and epiblast. A total of n = 15 embryos were imaged for both crosses. All scale bars 25 μm.

**Table S1.** Phase length.

| Embryo number | Mouse line | Phase length (mins) |  |  |
| --- | --- | --- | --- | --- |
|  |  | Pre-Migration/<br>Initiation | Migration | Post-Boundary |
| 1 | Lifeact-GFP_CD1 | - | 190 | 310 |
| 2 | Lifeact-GFP_CD1 | 215 | 135 | 100 |
| 3 | Lifeact-GFP_CD1 | 125 | 395 | 40 |
| 4 | Lifeact-GFP_CD1 | 60 | 165 | 270 |
| 5 | Lifeact-GFP_CD1 | 190 | 270 | 175 |
| 6 | Lifeact-GFP_CD1 | 155 | 235 | 225 |
| 7 | Lifeact-GFP_CD1 | 60 | 270 | - |
| 8 | Lifeact-GFP_CD1 | - | 110 | 220 |
| 9 | Lifeact-GFP_CD1 | - | 195 | 320 |
| 10 | Hex-GFP_mem-tdTom | 60 | 160 | 100 |
| 11 | Hex-GFP_mem-tdTom | 220 | 350 | 340 |
| 12 | Hex-GFP_mem-tdTom | 280 | 230 | 390 |
| 13 | Hex-GFP_mem-tdTom | 280 | 320 | 410 |
| 14 | Hex-GFP_mem-tdTom | 60 | 200 | 230 |
| 15 | Hex-GFP_mem-tdTom | 130 | 270 | 510 |
| 16 | Hex-GFP_mem-tdTom | 100 | 250 | 170 |
| 17 | Hex-GFP_mem-tdTom | 100 | 290 | - |
| 18 | Hex-GFP_mem-tdTom | - | 360 | - |
| 19 | Ttr-CRE: mem-tdTom | - | 200 | 100 |
| 20 | Ttr-CRE: mem-tdTom | - | 270 | 60 |
| 21 | Ttr-CRE: mem-tdTom | - | 170 | 180 |
|  | **total (mins)** | 2035 | 5035 | 4150 |
|  | **average (mins/embryo)** | 145.36 | 239.76 | 230.56 |

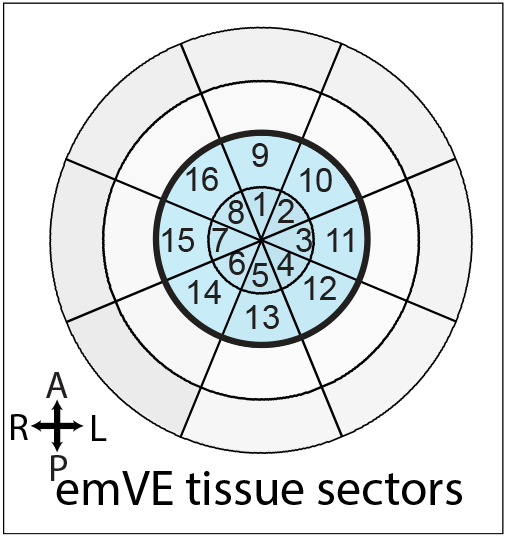

**Table S2.** Migration phase vector angles.

| Tissue sector | Rayleigh distribution | p-value |
| --- | --- | --- |
| 1 | 0.96 | 1.29E-08 |
| 2 | 0.93 | 2.78E-08 |
| 3 | 0.94 | 2.18E-08 |
| 4 | 0.96 | 1.08E-08 |
| 5 | 0.97 | 8.01E-09 |
| 6 | 0.96 | 1.25E-08 |
| 7 | 0.94 | 1.80E-08 |
| 8 | 0.95 | 1.68E-08 |
| 9 | 0.88 | 7.41E-08 |
| 10 | 0.76 | 7.52E-07 |
| 11 | 0.73 | 2.81E-06 |
| 12 | 0.81 | 2.75E-07 |
| 13 | 0.73 | 2.56E-06 |
| 14 | 0.64 | 8.36E-05 |
| 15 | 0.50 | 0.004423676 |
| 16 | 0.79 | 3.46E-07 |

